# PhysDock: A Physics-Guided All-Atom Diffusion Model for Protein-Ligand Complex Prediction

**DOI:** 10.1101/2025.04.28.650887

**Authors:** Kexin Zhang, Yuanyuan Ma, Jiale Yu, Huiting Luo, Jinyu Lin, Yifan Qin, Xiangcheng Li, Qian Jiang, Fang Bai, Jiayi Dou, Jie Zheng, Jingyi Yu, Liping Sun

## Abstract

Accurate prediction of protein-ligand complexes remains a central challenge in structural biology. Traditional methods are computationally inefficient and prone to local minima, whereas deep learning approaches struggle to capture structural flexibility and physical plausibility. We introduce PhysDock, a physics-guided diffusion model that uniquely integrates (i) all-atom diffusion to model ligand flexibility and protein precision-flexibility (*i*.*e*., subtle conformational adjustments); (ii) physical priors as diffusion conditioning, alongside two-phase physics guidance during the denoising diffusion to ensure physical plausibility. PhysDock demonstrates state-of-the-art performance in redocking benchmarks and excels in the more challenging cross-docking assessments. For practical utility, PhysDock (i) resolves cannabinoid receptor selectivity across diverse molecules, achieving accuracy comparable to experiments; (ii) distinguishes most drug candidates from weak binders in virtual screening of NTRK3 kinase, while uncovering novel candidates with structural insights. PhysDock serves as a versatile tool for protein-ligand complex prediction, with substantial potential to accelerate structure-based drug discovery.

## 1 Main

In structure-based drug discovery (SBDD), a pivotal task is to identify molecules that can modulate the biological functions of target proteins. This relies fundamentally on accurate structure prediction of protein-ligand complexes, including binding pockets, ligand orientations, protein side-chain conformations, and their interactions, all of which are governed by physics. First-principles methods, such as density functional theory, are theoretically accurate but computationally prohibitive^1,2^. Widely used tools, such as Glide^3^ and AutoDock Vina^4^, which rely on classical force fields and empirical scoring functions, respectively. Both are inefficient in exploring conformational landscapes and are prone to convergence at local minima, restricting their ability to find reliable structures^5–7^. These limitations compromise the accuracy of binding mode predictions, underscoring the need for more rigorous computational approaches.

Recent advances in deep learning (DL) have shown promises in protein-ligand complex prediction, including (i) SurfDock^8^, which incorporates a surface representation of the protein binding pocket to describe its geometry and complementarity, thereby enhancing prediction accuracy; (ii) Interformer^9^, which employs a graph transformer architecture with an interaction-aware mixture density network to capture binding affinity; (iii) Uni-Mol Docking V2^10^, which integrates pre-trained molecular and pocket encoders to ensure accurate geometric and chiral predictions; (iv) DiffDock^11,12^, which frames molecular docking as a generative modeling task, using a diffusion model over the non-Euclidean manifold of ligand poses to sample the best prediction; (v) DynamicBind^13^, which utilizes a geometric diffusion network to dynamically adapt protein conformations for optimal ligand binding. Additionally, large-scale generative models, such as AlphaFold3^14^, Chai-1^15^, and NeuralPLexer^16^, further improve the accuracy by predicting flexible structures for proteins and ligands using multimodal data representations and deeper neural networks.

However, critical challenges persist due to the lack of physical realism in the DL-based docking models. Many models are limited to semi-flexible docking, allowing ligand flexibility while treating protein structures as rigid. This simplification neglects the fundamental physical principle that both the ligand and protein undergo conformational changes upon binding. As a result, these models fail to capture the energetically favorable rearrangements of binding pocket residues, which are essential for accommodating diverse ligand scaffolds. Further limitations arise from physical implausibilities in modeling non-covalent interactions between proteins and ligands, as well as in the stereochemical integrity of ligands. For instance, key protein-ligand interactions (*e*.*g*., hydrogen bonds, *π*-stacking) are often poorly captured, even in models that achieve geometrically accurate ligand poses such as diffusion-based methods. Ligands may exhibit incorrect chirality or steric clashes resulting from distorted bond lengths and angles, even when their overall structure closely aligns with the ground truth. These limitations constrain their practical utility in drug discovery.

At the core of the issue is the absence of a mechanism to incorporate physical principles. Although physics-aware diffusion models have shown potential in motion generation^17^, 3D scene synthesis^18^, and protein structure generation^19^, such approaches have yet to be explored in the domain of molecular docking. PhysDock bridges this gap by integrating physics with the generative power of diffusion models, introducing key advances: (i) all-atom diffusion to model ligand flexibility and protein precision-flexibility—subtle conformational adjustments of binding pocket residues to adapt to varying binding modes; (ii) integration of physical priors as diffusion conditioning to enhance the physical plausibility of protein-ligand interactions, alongside a two-phase physics guidance during the denoising diffusion process leveraging pre-computed optimal conformer libraries and molecular mechanics force fields to correct chiral and steric errors in the ligands. Moreover, PhysDock employs an iterative sampling mechanism across distinct diffusion pathways, coupled with training-free clustering algorithms to systematically explore and identify optimal structures. Complemented by its streamlined architecture, PhysDock surpasses conventional generative models in sampling efficiency, making it well-suited for large-scale virtual screening applications.

PhysDock demonstrates dual capabilities in binding mode prediction: (i) without prior pocket information, it achieves the accuracy of large-scale generative models such as AlphaFold3 and Chai-1, while operating significantly faster; (ii) with prior pocket information, it establishes a new state-of-the-art (SOTA) performance across three redocking benchmarks, PoseBusters^20^, DeepDockingDare, and our curated PhiBench dataset. Specifically, PhysDock achieves 5%-10% higher success rates than current best docking models, and recovering protein-ligand interactions comparable to semi-flexible docking using rigid ground-truth pockets. Notably, in the more challenging cross-docking assessments involving protein structures resolved in complex with different ligands, PhysDock outperforms leading docking models, achieving > 15% improvement over leading docking models and the highest recovery rates of protein-ligand interactions. To demonstrate its practical utility in drug discovery applications, we first employ PhysDock to predict CB1 versus CB2 receptor binding selectivity across a diverse set of ligands, achieving results comparable to experimental measurements. We then implement a virtual screening pipeline targeting NTRK3 kinase based on PhysDock, effectively distinguishing most high-affinity hits, including drug candidates, from low-affinity binders while also predicting novel candidate molecules with structural insights. These results underscore PhysDock’s potential to deliver previously unattainable accuracy and efficiency in protein-ligand complex prediction.

## 2 Results

### 2.1 PhysDock overview

PhysDock is a physics-guided all-atom diffusion model designed for protein–ligand complex prediction (Fig. 1a, Methods, SI Text 1-4). While building upon broader advances in AlphaFold2^21^, AlphaFold3^14^, Llama3^22^, and Stable Diffusion 3^23^, PhysDock introduces critical innovations specifically tailored for molecular docking. Uniquely, it integrates physical priors on protein-ligand interactions as diffusion conditioning, alongside a two-phase physics guidance during the denoising process leveraging pre-computed optimal conformer libraries and molecular mechanics force fields to enforce physical plausibility. At its core, PhysDock employs all-atom diffusion to simultaneously generate 3D coordinates of both the protein and ligand, explicitly modeling ligand flexibility and protein precision-flexibility (*i*.*e*., subtle conformational adjustments that accommodate diverse binding modes). This design embraces the fundamental physical principle of mutual induced fit upon binding. Trained on Plinder^24^, a high-quality, well-annotated dataset curated for molecular docking tasks, the model consists of two primary modules: DiffusionConditioning and DiffusionModule.

**Figure 1.**
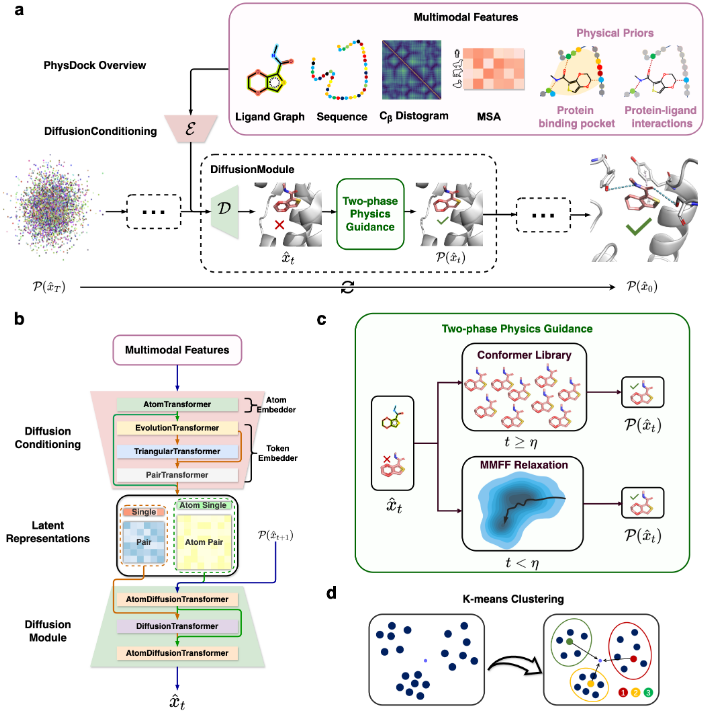
(a) Overview of PhysDock’s architecture. The multimodal features, including ligand graphs, protein sequence features, backbone C_*β*_ distance-gram, MSA features and physical priors on protein-ligand interactions, are encoded by the DiffusionConditioning module. Subsequently, in each denoising step within the DiffusionModule, represented by the dashed box, the denoised molecule structure undergoes a two-phase physics guidance to enforce physical plausibility. After *T* sampling steps, PhysDock generates protein-ligand complex structures from pure noise. (b) Network architecture of PhysDock. (c) Two-phase physics guidance in stepwise denoising. The denoised ligand structure is projected using either a pre-computed conformer library or molecular mechanics force field relaxation determined by an adaptive factor *η*. (d) K-means clustering to identify optimal representative structures. All sampled complexes are clustered based on PAL-RMSD as the distance metric between sample points, with the sample closest to the center of each cluster is selected as the representative structure. These representatives are then ranked based on their proximity to the global centroid of all samples.

The DiffusionConditioning module processes multimodal input features (Fig. 1a, SI Table 1), including ligand molecule graph (*i*.*e*., atomic-scale chemical and geometrical features of molecules), protein sequence, backbone C_*β*_ distance diagram, multiple sequence alignments (MSA), and physical priors. Notably, the physical priors comprise protein binding pocket residues and key residues involved in various types of non-covalent interactions with the ligand, thereby enhancing the physical plausibility of protein–ligand interactions. To improve robustness and generalizability, PhysDock incorporates a wide range of data augmentation strategies, such as spatial cropping, MSA sampling, and probabilistic activation of physical priors. For instance, 50% of randomly selected key residues have a 50% chance of being included in each diffusion process. These features are then encoded into atom- and token-level latent space representations (with each token corresponding to a protein residue or ligand atom), which serve as denoising conditions for the subsequent diffusion process (Fig. 1b).

The DiffusionModule is a dual-scale diffusion transformer (Fig. 1b). It operates at atomic resolution utilizing a full-atom diffusion transformer without any masking to model attention across all pairs of geometrically neighboring atoms, while incorporating a token-based processing block in its intermediate layers through a token-level diffusion transformer to capture residue-level interactions. This hierarchical feature encoding enables multiscale modeling of protein-ligand complexes—from fine-grained atomic details to broader topological constraints. Notably, PhysDock applies a two-phase physics guidance to correct chiral and steric errors in ligands. In each denoising diffusion step, PhysDock projects the denoised ligand molecules into a physically plausible conformational space, guiding the subsequent denoising diffusion step. The projection strategy varies depending on the noise level and an adaptive factor *η*, which is dynamically adjusted based on the final sampling results of each round. In high-noise diffusion steps (*t* ≥*η*), PhysDock employs an exploring-matching-aligning-replacing strategy to correct chiral and steric errors in ligand molecules, which explores the pre-computed conformer library to match the closest conformation to the denoised ligand based on atom distance matrices, followed by alignment and replacement. In low-noise diffusion steps (*t* < *η*), PhysDock adopts a relaxing-aligning-replacing strategy to further correct steric clashes in ligand molecules, performing a few steps of relaxation based on molecular mechanics force fields (MMFF) to optimize the denoised ligand, followed by alignment and replacement.

PhysDock implements an iterative sampling strategy to thoroughly explore the conformational landscape and mitigate convergence to local optima^25,26^. This is accomplished through multiple rounds of Diffusion Conditioning and Physics Guided Sampling Diffusion, repeated until a desired number of samples are obtained. Importantly, each iteration incorporates MSA resampling to generate diverse latent-space representations, significantly improving the efficiency of conformational space exploration^27–29^. At this stage, PhysDock generates a substantial number of physically plausible and diverse protein-ligand complexes along distinct diffusion pathways through iterative sampling while remaining close to the data distribution in a holistic manner. Subsequently, K-means clustering^30^ is employed to identify optimal representative structures (Fig. 1d), a method proven effective in the conformational analysis of proteins^31^, nucleic acids^26^, and small molecules^32^. The top-5 representative structures are selected as the nearest neighbors to the centers of five distinct clusters and ranked according to their proximity to the global centroid of all samples. As expected, the docking success rate is positively correlated with the number of samples generated during inference (Extended Data Fig. 1a). This ranking strategy effectively improves structure prediction accuracy by identifying hidden intrinsic structural patterns, outperforming neural ranking scores, such as the ConfidenceModule in AlphaFold3 (Extended Data Fig. 1bc). It also captures structural diversity among predicted samples, potentially approximating a Boltzmann-like distribution given sufficient sampling (SI Figure 14).

Additionally, PhysDock exhibits significantly enhanced efficiency compared to large-scale generative models due to its streamlined architecture, including utilizing downscaled deep networks like PairFormer, removing the recycling process in the trunk module, and decreasing the number of diffusion steps during sampling. As a result, the compile-free inference time of PhysDock achieves a 4-16 times faster than compile-free Chai-1 and 2-4 times faster than compiled AlphaFold3 across token sizes (Extended Data Table 1). When binding pocket information is provided, PhysDock can be further accelerated by region-specific cropping, which significantly reduces the sampling space. Together, these advances enable PhysDock to predict protein-ligand complexes with high accuracy, physical plausibility, and efficiency, making it a powerful tool for structure-based drug design. The data and code for PhysDock are publicly available on open-source platforms.

### 2.2 PhysDock achieves SOTA performance in redocking benchmarks

To evaluate the robustness and generalizability, we conducted a comprehensive benchmarking study against available docking methods: (i) classical docking tools, *i*.*e*., AutoDock Vina^4^, Glide^3^; (ii) semi-flexible DL models, *i*.*e*., SurfDock^8^, Interformer^9^, Diffdock^11^, Diffdock-L^12^, Uni-mol Docking V2^10^; (iii) flexible DL models, *i*.*e*., AlphaFold3^14^, Chai-1^15^, DynamicBind^13^, NeuralPLexer^16^ (SI Text 5). The evaluation was performed using three datasets: PoseBusters^20^, DeepDockingDare (DDD, https://github.com/DSDD-UCPH/DeepDockingDare, and our curated PhiBench dataset. The PoseBusters dataset, comprising 428 protein-ligand complexes, is the current gold standard to evaluate the accuracy of ligand binding modes predicted by docking methods. It provides a metric system, PB-valid, encompassing 18 criteria to assess the chemical and geometric consistency of ligands. The DDD dataset comprises 425 recently resolved, structurally diverse protein-ligand complexes with substantial coverage of membrane proteins (38%), specifically G-protein-coupled receptor (GPCR)–G protein complexes (23%). Our curated PhiBench dataset comprises 206 newly resolved protein-ligand complexes, deposited between June and December 2024, and selected to ensure sufficient sequence diversity (SI Figure 15).

All models were evaluated using the protein-aligned ligand root-mean-square deviation (PAL-RMSD) in two scenarios: without and with prior information about the protein binding pocket. For each protein-ligand complex, 40 candidate structures were generated and ranked using method-specific scoring metrics (*e*.*g*., binding free energy for Glide, clustering-based scores for PhysDock, and confidence scores for other deep learning models). Two docking success rates were computed as having at least one top-5 ranked pose satisfying either of the following criteria: (1) PAL-RMSD < 2Å, or (2) PB-valid & PAL-RMSD < 2Å. In docking tasks without prior binding pocket information (Fig. 2a, upper panel), under the PAL-RMSD < 2Å criterion, PhysDock achieves a docking success rate of 65.2% on PoseBusters, 53.5% on DDD, and 61.2% on PhiBench. Under the stricter criterion of PB-valid & PAL-RMSD < 2Å, the docking success rates are 61.3% on PoseBusters, 49.3% on DDD, and 49.5% on PhiBench. This performance is comparable to that of the leading large-scale generative models, achieving slightly lower success rates than AlphaFold3 on PoseBusters and DDD, but slightly outperforming AlphaFold3 on the more recently curated PhiBench dataset. Additionally, it achieves similar success rates to Chai-1 on PoseBusters and surpasses Chai-1 on both DDD and PhiBench. Plausibly, PhysDock delivers this remarkable performance while offering a 2–4 times speedup over AlphaFold3 and a 4–16 times speedup over Chai-1 across different token sizes (Extended Data Table 1).

**Figure 2.**
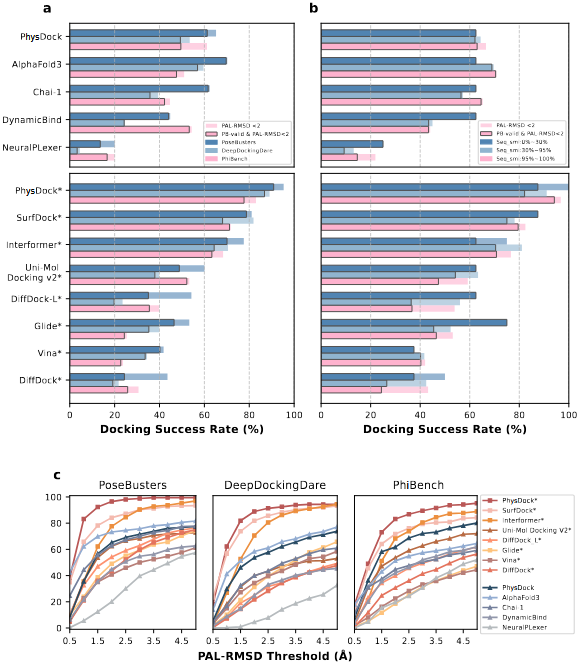
Performance of various docking models in redocking benchmarks, with and without prior pocket information. **(a)** Docking success rates of different models across the PoseBusters dataset (N=428), the DeepDockingDare dataset (N=356) and PhiBench dataset (N=206). Asterisks (*) denote predictions with prior pocket information. **(b)** Docking success rates of different models on the PoseBusters dataset, categorized by sequence similarity to the plinder training set (PoseBusters: 428 samples, subsets: 8/112/308 for low/medium/high similarity). **(c)** Docking success rate of different models under increasing PAL-RMSD thresholds (from 0.5 to 5.0Å).

In docking tasks with prior binding pocket information (Fig. 2a, lower panel, SI Table 2), PhysDock establishes SOTA performance across all benchmarks. It achieves a PAL-RMSD < 2Å success rate of 95.3% on PoseBusters, 89.1% on DDD, and 83.0% on PhiBench. Notably, PhysDock demonstrates superior performance compared to the reported, but non-open-source, docking mode of AlphaFold3 (90.2% success rate on PoseBusters) and Chai-1 (81.2% success rate on PoseBusters). Under the stricter criterion of PB-valid & PAL-RMSD < 2Å, PhysDock maintains SOTA performance with a top-5 success rate of 90.9% on PoseBusters, 87.0% on DDD, and 77.7% on PhiBench. As a result, PhysDock outperforms best docking baselines by approximately 5%-10% under both criteria. A detailed analysis of the 18 criteria comprising the PB-valid metric reveals improved docking success rates when either prior binding pocket information is provided or physics guidance is applied (Extended Data Fig. 2). Notably, marked improvements are seen in the tetrahedral chirality and minimum distance to protein checks, underscoring the effectiveness of incorporating physics into the docking process. These findings highlight PhysDock’s unique strength in enhancing the docking accuracy by integrating physical priors and physics guidance within the all-atom diffusion model. In contrast, classical docking tools like Glide (34.4% success rate) and AutoDock Vina (33.6%) show significantly lower performance on the PoseBusters benchmark compared to DL-based approaches, further emphasizing the advancements enabled by modern generative methods.

Next, we conducted an extended evaluation using the PoseBusters benchmark, categorizing test cases based on their sequence similarity relative to our training dataset, Plinder^24^. Without prior binding pocket information (Fig. 2b, upper panel), PhysDock achieves performance comparable to AlphaFold3 at low (0% ∼30%) sequence similarity, and shows slightly lower success rates at medium (30% ∼95%) and high (95% ∼100%) sequence similarity levels. With prior binding pocket information (Fig. 2b, lower panel), PhysDock achieves SOTA performance across all sequence similarity categories and, more importantly, maintains consistent success rates across these categories for all three benchmarks (SI figure 16). This consistency not only confirms the absence of training data leakage but also demonstrates PhysDock’s unique ability to capture intrinsic protein-ligand interaction patterns, rather than relying solely on sequence homology, further enhancing its robustness. Finally, we analyzed the docking success rates of available methods across all three benchmarks using multiple PAL-RMSD thresholds (0.5 ∼ 5Å) (Fig. 2c). Without prior binding pocket information, PhysDock demonstrates performance on average comparable to AlphaFold3. With prior binding pocket information, PhysDock achieves higher success rates at lower PAL-RMSD thresholds (*e*.*g*., 0.5 to 1.5Å), underscoring its superior ability to generate high-precision binding poses. Together, these results underscore PhysDock’s accuracy and robustness in different redocking scenarios.

Beyond evaluating the geometric resemblance between predicted and ground-truth structures, We conducted a comprehensive analysis of the protein–ligand interaction recovery rate (IRR) to quantify how effectively each docking method reproduces these interactions across all three benchmark datasets (Fig. 3a-c, SI Table 3). IRR is computed as the fraction of non-covalent interactions in the experimentally resolved structures that are successfully predicted by the model, including hydrogen bonds, hydrophobic interactions, salt bridges, *π*-stacking, and *π*-cation interactions. These interactions are identified using the Protein-Ligand Interaction Profiler (PLIP)^33^. With prior binding pocket information, PhysDock outperforms all three flexible docking models, *i*.*e*., DynamicBind, AlphaFold3, and Chai-1, across all types of non-covalent interactions. It demonstrates comparable performance to the best semi-flexible docking methods, where the ground-truth binding pocket residues, including side chains, are given and held rigid. Averaged across the three datasets, PhysDock recovers 50.3% of hydrogen bonds and 62.8% of hydrophobic interactions—two of the most frequently occurring interaction types—compared to Uni-Mol Docking V2, which recovers 62.2% and 52.5%, and SurfDock, which recovers 59.5% and 55.1%, respectively.

**Figure 3.**
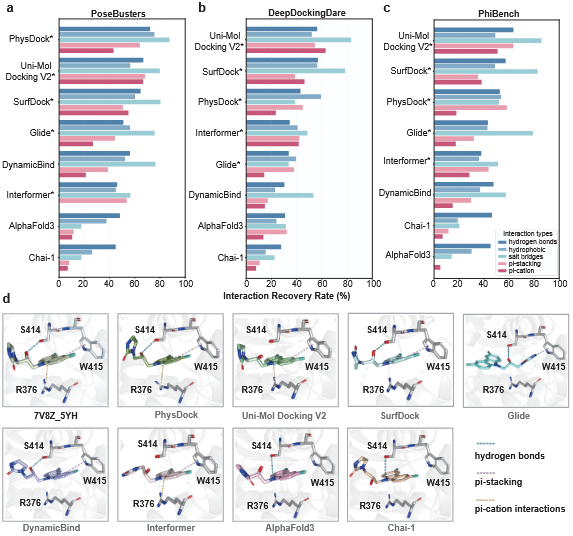
Interaction Recovery Rates (IRR) and predicted binding modes of various docking models in redocking benchmarks, with and without prior pocket information. (a-c) IRR (%) for different types of non-covalent interactions across three datasets: PoseBusters, DeepDockingDare, and PhiBench. Asterisks (*) denote predictions with prior pocket information. (d) Predicted binding modes from different models, including PhysDock, Uni-Mol Docking V2, SurfDock, Glide, DynamicBind, Interformer, AlphaFold3, and Chai-1, compared with the experimental structure (PDB ID: 7V8Z_5YH). Dashed lines indicate key interactions: hydrogen bonds (blue), *π*-stacking (pink), and *π*-cation interactions (yellow). Hydrophobic interactions are omitted for clarity.

Fig. 3d presents a case study from the PoseBusters dataset. The experimental structure (PDB ID: 7V8Z) features four non-covalent interactions: (i) one hydrogen bond between the hydroxyl group of S414 and the oxygen atom of the ligand (CCD ID: 5YH), (ii) one *π*-stacking between the indole ring of W415 and the ligand, (iii) one *π*-cation interaction between the guanidinium group of R376 and the ligand’s benzene ring and several hydrophobic interactions that are not explicitly shown in the figure for clarity. While all docking methods except Glide predict approximately correct ligand poses in the binding pocket (PAL-RMSD of approximately 2.4Å), PhysDock uniquely recovers all three key interactions. In contrast, other methods either partially recover the interactions or introduce incorrect interaction sites. For example, AlphaFold3 predicts a hydrogen bond between S414 and a different atom in the ligand, which is not observed in the ground-truth structure. The high recovery rate of protein-ligand interactions underscores PhysDock’s effectiveness in capturing physically plausible non-covalent interactions, validating the integration of physical priors and physics guidance in its design.

In summary, PhysDock marks a significant step forward in robustly predicting accurate and physically plausible protein–ligand complex structures, while effectively capturing key non-covalent interactions. Its strong performance underscores its potential for large-scale docking applications, where both reliable ligand poses and interaction fidelity are critical for ranking candidate binding modes.

### 2.3 PhysDock demonstrates superior cross-docking accuracy with varying ligands

To evaluate the practical utility of DL-based models in real-world docking tasks, we curated a time-split cross-docking test set consisting of 25 proteins, each in complex with two different ligands—one resolved before June 2024 and the other after, to avoid potential overlap with the training data. This resulted in a total of 50 test cases, each involving either protein_1_-ligand_2_ or protein_2_-ligand_1_, where subscripts 1 and 2 denote distinct experimental structures differing in protein binding pocket conformations, ligand identity, and their interactions. In docking cases where ligand_2_ was docked to protein_1_, the binding pocket residues of protein_1_ were provided as input, and the protein_2_–ligand_2_ complex structure was used as the ground truth. This cross-docking test set allows us to evaluate: (i) the prediction of new ligand binding modes within a prior binding pocket resolved with another ligand; and (ii) the stability of model performance in different resolved structures of the same protein with potential compositional or conformational variations of the binding pocket. We evaluated five top-performing docking models from redocking benchmarks—PhysDock, AlphaFold3, Interformer, Uni-Mol Docking V2, and SurfDock.

We first computed the docking success rates of different models under two thresholds: PAL-RMSD < 2Å and PB-valid & PAL-RMSD < 2Å (Fig. 4a). PhysDock surpasses other docking models by at least 15% under both criteria and maintains high success rates even at lower PAL-RMSD thresholds (*e*.*g*., 0.5 to 1.5Å), underscoring its significant advantage in cross-docking tasks. We then quantified the interaction recovery rates (IRR) of different models for non-covalent protein-ligand interactions (Fig. 4b). PhysDock outperforms other docking models in recovering all five types of interactions, with particularly strong performance in recovering salt bridge interactions. Together, these results demonstrate PhysDock’s superior performance over current top-performing docking models in both geometric accuracy and physical plausibility on challenging cross-docking tasks. This advantage arises from PhysDock’s integration of physical priors and physics guidance into an all-atom diffusion framework, which directs the binding pocket to adapt its conformation to diverse ligand binding modes.

**Figure 4.**
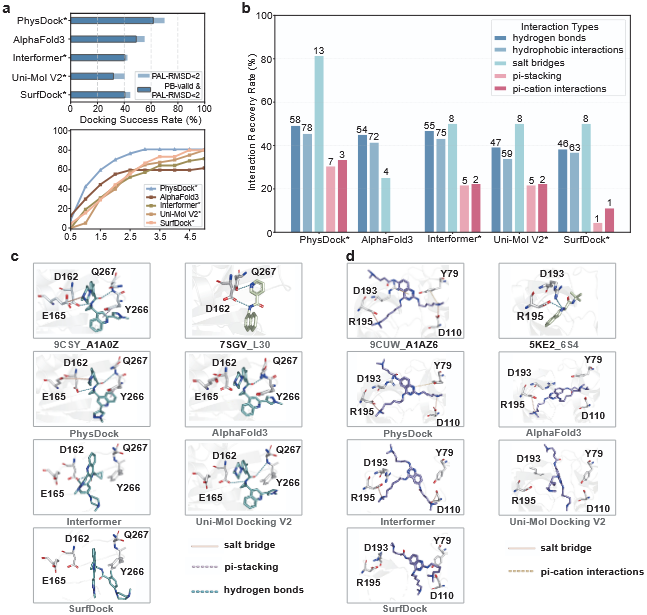
Comparative analysis of various docking models in cross-docking assessments. **(a)** Docking success rates of different models under two criteria: PAL-RMSD < 2Å and PB-valid & PAL-RMSD < 2Å. **(b)** Interaction Recovery Rate (IRR) for five types of non-covalent interactions across four models. **(c-d)** Two representative cross-docking cases comparing the predicted binding modes from different models with the ground truth. Dashed lines indicate key interactions: salt bridges (brown), hydrogen bonds (blue), *π*-stacking (pink), and *π*-cation interactions (yellow). Hydrophobic interactions are omitted for clarity.

Fig. 4c-d illustrate two representative cases from the cross-docking test set. The first case involves docking the ligand from PDB ID 9CSY to the protein from PDB ID 7SGV, which features differing side-chain conformations in the binding pocket (Fig. 4c). Among the docking models, PhysDock, AlphaFold3, and Uni-Mol Docking V2 predict reliable ligand poses within the binding pockets. Notably, PhysDock recapitulates the key protein-ligand interactions observed in the ground-truth structure, while other models only partially recover them. In the reverse docking case (Extended Data Fig. 3a), where the ligand from PDB ID: 7SGV is docked to the protein from PDB ID: 9CSY, PhysDock again produces a reliable binding pose and successfully recovers one of the two hydrogen bonds with the pocket residues, while other models fail to recover either the ligand pose or key interactions. The second case involves docking the ligand from PDB ID: 9CUO to the protein from PDB ID: 7BQV, representing a more challenging cross-docking task due to substantial differences in the binding pocket conformations, ligand poses, and protein-ligand interaction patterns (Fig. 4d). PhysDock and Interformer predict binding poses with relatively small deviations from the ground truth and partially recover the protein-ligand interactions. In contrast, other models show larger deviations, with SurfDock producing molecular overlaps within the binding pocket, resulting in physically incorrect ligand conformations. In the reverse docking case (Extended Data Fig. 3b), where the ligand from PDB ID: 5KE2 is docked to the protein (PDB ID: 9CUW), PhysDock, Interformer, and SurfDock are the only models that generate approximately reliable binding poses and partially recover key residues involved in non-covalent interactions with the ligand. We present an additional cross-docking case between PDB ID: 9CUO and PDB ID: 7BQV (Extended Data Fig. 3c-d), a task that poses significant challenges due to an additional sequence segment in the binding pocket of 7BQV not present in 9CUO. PhysDock uniquely achieves ligand binding pose close to the ground truth by repositioning the additional sequence segment to accommodate the different binding mode. Other models, however, confine ligands to predefined binding pockets, resulting in clashes, while AlphaFold3, despite supporting flexible docking, remains limited by its MSA-derived pocket predictions. These cross-docking tasks further highlight PhysDock’s superior ability to predict both ligand binding poses and protein-ligand interactions based on protein structures bound to different ligands, closely mirroring real-world docking challenges aimed at discovering novel ligands.

### 2.4 PhysDock resolves cannabinoid receptor selectivity with structural insights

We applied PhysDock to predict receptor selectivity between the closely related CB1 and CB2 cannabinoid receptors across diverse ligands, addressing a longstanding challenge in GPCR drug discovery. Despite sharing 44% sequence identity and significant structural similarity, these receptors mediate distinct physiological pathways: CB1 primarily regulates central nervous system functions, with its antagonists showing therapeutic potential for metabolic disorders^34,35^, while CB2 modulates peripheral immune responses, making its agonists promising candidates for the treatment of inflammatory and neuropathic pain^36,37^. Using cryo-EM structures of active CB1 (PDB: 6KPG) and CB2 (PDB: 6KPF) in complex with G proteins^38^, we first perform a comparative analysis of their binding modes and protein-ligand interaction profiles (Fig. 5a). Note that the two ligands share a similar scaffold, with the CB1-bound ligand featuring a more extended carbon tail compared to the CB2-bound ligand. Despite their nearly identical binding pockets and binding poses, CB1 forms eight hydrophobic contacts with its ligand, while CB2 establishes a more elaborate interaction profile featuring eleven hydrophobic contacts plus one hydrogen bond with its ligand. These distinct binding modes provide a structural basis for the observed ligand selectivity differences. We first performed redocking for the two cannabinoid receptor complexes using PhysDock. In addition to accurately predicting the binding poses, PhysDock successfully recovers most protein-ligand interactions. Specifically, 7 out of 8 hydrophobic interactions are recovered for the CB1 receptor, and 1 out of 1 hydrogen bond and 6 out of 11 hydrophobic interactions are recovered for the CB2 receptor (Extended Data Fig. 4). We then selected 40 diverse compounds whose experimental binding free energies for the CB1 and CB2 receptors were calculated based on their experimentally measured inhibition constants, *K*_*i*_, as follows:

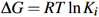

where *R* is the gas constant (1.987 × 10 ^−3^ kcal mol^−1^ ·K), and *T* denotes room temperature (298 K). For cases where binding energies approached instrumental detection limits (*e*.*g*., > 30000 nM for CB1; > 10000 nM for CB2), these values were used for the subsequent analysis.

**Figure 5.**
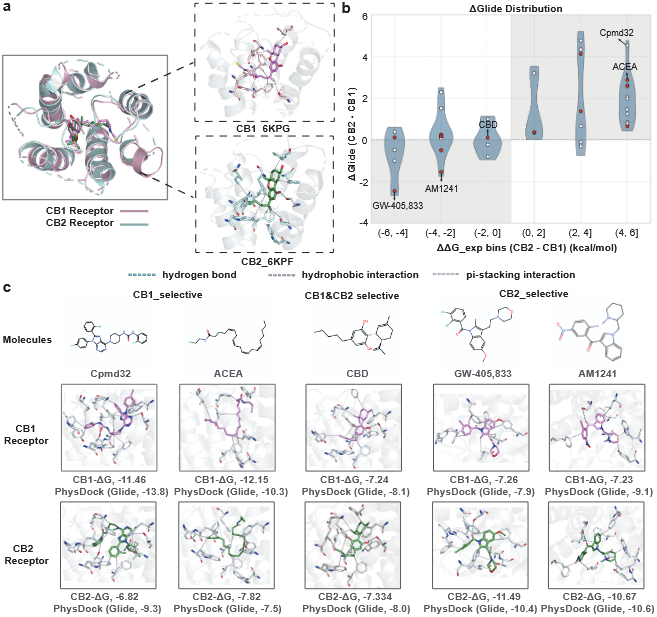
Comparative analysis of cannabinoid receptor selectivity across diverse compounds. **(a)** Overlay of CB1 (PDB: 6KPG) and CB2 (PDB: 6KPF) receptor structures showing their ground-truth ligand binding modes, with key interacting residues highlighted. **(b)** Glide score difference for all compounds docked to CB1 and CB2 receptors using PhysDock (ΔGlide = CB2 − CB1) plotted against their experimental binding free energy difference (ΔΔ*G*_exp_), with representative drug molecules highlighted in red. **(c)** Docking poses and 2D structures of six representative compounds bound to CB1 and CB2 receptors, with key interactions labeled.

We performed molecular docking of these compounds to CB1 and CB2 cryo-EM structures using PhysDock, with subsequent binding free energy estimation using Schrödinger’s Glide scoring function^39^. While Glide scores are known to exhibit size-dependent bias (*i*.*e*., favoring larger molecules capable of more interactions, SI Figure 17), this systematic error is expected to have a negligible impact when evaluating relative receptor selectivity of the same ligand. Remarkably, PhysDock successfully distinguished receptor-selective molecules compared to their experimental binding free energies, with the majority of CB1- and CB2-selective ligands correctly classified in their respective quadrants (Fig. 5b, Supplementary Table 4). Fig. 5c illustrates representative molecules spanning different selectivity profiles: CB1-selective (compound 32 and ACEA), non-selective (CBD), and CB2-selective (GW-405,833 and AM1241). Notably, all except compound 32, are drug candidates currently in preclinical development or clinical trials. Although the two CB1-selective ligands maintain generally comparable binding poses in both receptors, they adopt a slightly strained geometry in CB2 compared to CB1, suggesting a less favorable fit. In contrast, the non-selective ligand adopts nearly identical poses in both CB1 and CB2 receptors, preserving similar hydrophobic interaction networks and forming the same hydrogen bond. The two CB2-selective ligands also display comparable poses but engage in more extensive hydrophobic interactions in CB2 than in CB1, likely explaining their higher predicted binding affinities. These distinct interaction patterns reflect receptor-specific non-covalent interactions that underlie the observed differences in their binding affinity. Collectively, these findings highlight PhysDock’s utility in identifying receptor-selective drug candidates within structure-based drug discovery pipelines.

### 2.5 PhysDock separates high- and low-affinity NTRK3 ligands in virtual screening

Leveraging PhysDock, we implemented a virtual screening pipeline targeting the NTRK3 kinase. As a high-affinity receptor for neurotrophin-3 (NT3), NTRK3 regulates neural development and function^40^. Chromosomal translocations involving NTR can generate fusion proteins, most notably ETV6–NTRK3 (EN), whose constitutive activation leads to aberrant kinase activity associated with various cancers^41^. Therefore, identifying molecules that inhibit NTRK3 holds significant therapeutic potential. We performed virtual screening against NTRK3 kinase using its crystal structure (PDB: 6KZD)^42^ and the SPECS compound database (Fig. 6a). Our pipeline comprised three steps: (i) molecular docking of all database compounds using PhysDock to generate optimal protein-ligand complexes; (ii) 100-step side-chain relaxation of the binding pocket using MMFF optimization; and (iii) affinity-based ranking of all complexes using Glide scores to classify high-affinity hits and low-affinity binders. In addition, we screened 15 drug candidates with low IC_50_ values, which are under varying development phases, from preclinical compounds (*e*.*g*., PROTAC CPDs^43^, CH7057288^44^, Altiratinib^45^), to clinical trial molecules (*e*.*g*., Lestaurtinib^46^ and Selitrectinib^47^), and to approved drugs (*e*.*g*., Repotrectinib^48^, Entrectinib^49^, and Larotrectinib^50^) (SI Table 5). To further assess PhysDock’s discrimination capability, we also screened 59 known low-affinity NTRK3 binders (IC_50_ > 1 *μ*M, > 10 *μ*M, or not measurable). These compounds were strategically derived from known drug candidates through targeted modifications (*e*.*g*., benzene ring substitutions), preserving the core scaffold while largely reducing binding affinity—thus serving as challenging negative controls.

**Figure 6.**
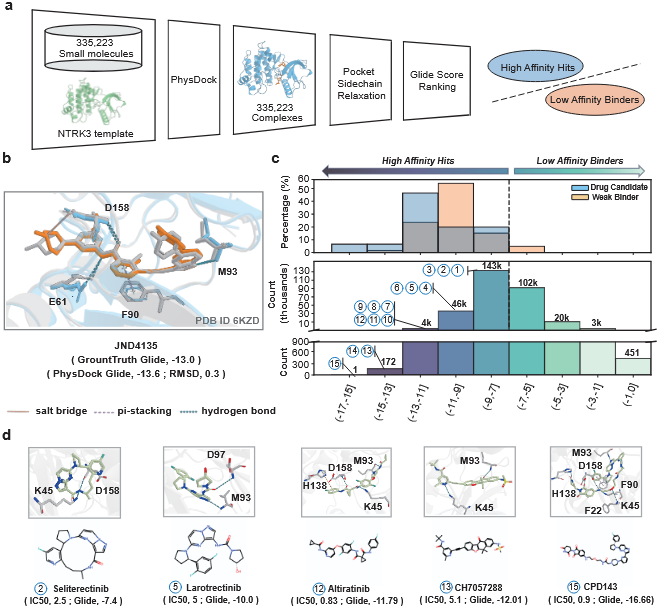
PhysDock effectively distinguishes most known drug candidates from weak binders of the NTRK3 kinase. **(a)** Virtual screening workflow for NTRK3 kinase using PhysDock, which includes small molecules from the SPECS database, 15 known drug candidates, and 59 weak binders. Pocket sidechain relaxation was performed for 100 steps prior to Glide score ranking. **(b)** Overlay of binding modes predicted by PhysDock (orange) and the ground truth (grey, PDB ID: 6KZD), with key interacting residues highlighted. Hydrophobic interactions are omitted for clarity. **(c)** Glide score distribution from PhysDock screening of NTRK3 kinase-targeted small molecules, classifying compounds as high-affinity hits or low-affinity binders. **(d)** Docking poses of known NTRK3 kinase inhibitors, with key interacting residues highlighted. Corresponding 2D structures are displayed below.

We first evaluated the predicted binding mode of JND4135 using PhysDock and found strong agreement with the experimentally determined crystal structure (Fig. 6b). While PhysDock does not recover the *π*-stacking and predicts an additional salt bridge, it accurately recapitulates the ligand binding pose and preserves all three hydrogen bonds present in the crystal structure. The predicted complex structure exhibits excellent geometric fidelity (PAL-RMSD = 0.3Å) and nearly identical Glide score (−13.6 kcal mol^−1^) to the ground truth (−13 kcal mol^−1^), highlighting PhysDock’s ability in generating both structurally and energetically reliable binding modes. Notably, kinases such as NTRK3 possess highly druggable binding pockets, likely due to their characteristic solvent-exposed polar front pockets and hydrophobic back cavities. As expected, more than half of the screened molecules (193K) are identified as high-affinity hits with Glide scores below –7 kcal mol^−1^, while the remaining (125K) are classified as low-affinity binders with scores above –7 kcal mol^−1^ (Fig. 6c). While Glide scoring tends to favor larger molecules, as seen in the screening results from the SPECS database, this size-related bias is not seen among the drug candidates and known weak binders (SI Figure 18). As expected, all drug candidates and most weak binders exhibit high-affinity predictions (Glide scores < –7 kcal mol^−1^), reflecting their shared score scaffolds. Note that PROTAC compounds show relatively stronger binding affinities (−11 to −17 kcal mol^−1^), likely attributable to their larger molecular size and bivalent binding pose. Despite some overlap, PhysDock distinguishes most known drug candidates from weak binders, as evidenced by their shift toward lower Glide scores (Fig. 6c; SI Table 6). These findings demonstrate PhysDock’s ability to effectively differentiate strong binders from weak ones.

Finally, we present the binding modes of both known drug candidates targeting NTRK3 kinase and newly identified high-affinity hits from virtual screening across varying Glide score ranges (Fig. 6d, Extended Data Fig. 5, Extended Data Fig. 6). The predicted complexes of drug candidates partially recapitulate key hydrogen bond or *π*-stacking interactions observed in the crystal structure, most commonly involving pocket residue M93, and in some cases D158 and/or F90. Similarly, the identified novel compounds, despite their diverse scaffolds, retain subsets of these critical interactions with pocket residues, suggesting strong inhibitory potential. Notably, the predicted complex with compound 2ADB96 (Glide score: –12.2 kcal mol^−1^) recapitalizes all four key interactions observed in the crystal structure (Extended Data Fig. 6b). Collectively, these results illustrate the effectiveness of PhysDock in identifying high-affinity candidate compounds targeting NTRK3 kinase and underscore its practical utility in real-world virtual screening.

## 3 Discussion

PhysDock overcomes current limitations in accuracy and physical plausibility in classical and DL-based docking methods. It is a physics-guided all-atom diffusion model to enable ligand flexibility and protein precision-flexibility, which incorporates DiffusionConditioning to incorporate physical priors on binding pockets and protein-ligand interactions, and SampleDiffusion to utilize pre-computed optimal conformer library and molecular mechanics force fields to correct ligand chiral and steric errors. Moreover, its iterative sampling strategy and clustering algorithms for ranking enable the thorough exploration and subsequent identification of accurate and diverse protein-ligand complex structures. Finally, its specific model architecture significantly enhances efficiency, achieving a 2-16 times speedup over large-scale generative docking models, making it well-suited for large-scale docking tasks. PhysDock’s capabilities are demonstrated in both redocking and cross-docking benchmarks, where it outperforms current docking methods in docking success rates and the recovery of protein-ligand interactions. Specifically, it achieves state-of-the-art performance on redocking benchmarks, outperforming the best open-source docking models by 5%-10% in success rate with prior pocket information. It also excels in the more challenging cross-docking assessments involving protein structures resolved with varying ligands, surpassing leading docking models by over 15% in success rate. For practical utility, PhysDock was successfully applied to resolve cannabinoid receptor selectivity across diverse molecules, and to the virtual screening of the NTRK3 kinase using the SPECS library of over 335,000 molecules, separating existing drug candidates at various stages of development from low-affinity binders and uncovering novel candidate molecules with structural insights. Together, PhysDock enables accurate, physically plausible, and efficient protein–ligand complex prediction, driving progress in structure-based drug discovery.

Despite its strengths, several aspects of PhysDock require improvement to further enhance its applicability in structure-based drug design (SBDD). First, PhysDock is currently optimized for predicting interfaces between proteins and single small-molecule ligands, limiting its applicability to more complex interactions involving peptides, nucleic acids, or multiple co-binding ligands. To address this limitation, we plan to expand the training data to include such diverse protein–ligand complexes. Second, the model’s sensitivity to backbone dynamics is limited, restricting its ability to perform docking on apo protein structures. We shall mitigate this issue by including apo structures, leveraging synthetic data from molecular dynamics simulations, and applying various data augmentation techniques. Third, PhysDock’s two-phase physical projection strategy is currently applied only to ligand molecules, excluding protein binding pocket residues to preserve sampling efficiency. However, this simplification may limit the physical plausibility of predicted protein–ligand interactions. To address this, we plan to integrate GPU-accelerated or deep learning–based atomistic force fields to improve efficiency and enable the inclusion of binding pocket residues in the physical projection process. Last, while PhysDock ensures the stereochemistry of small molecules through physical projection and iterative sampling, the network itself remains insensitive to stereochemical attributes such as chirality. This could be addressed by incorporating physics-informed neural network architectures and physics-aware criteria. These improvements are expected to further strengthen PhysDock’s capabilities, facilitating more accurate and reliable predictions in protein-ligand interactions.

In summary, accurate protein–ligand complex prediction holds transformative potential for both structural biology and therapeutic development. By integrating physics with the generative power of diffusion models, PhysDock offers a robust platform for structure-based drug design. With continued development, it is poised to become a valuable tool for discovering and optimizing novel ligands for both fundamental research and drug discovery applications.

## 4 Methods

### Data collection

The training and validation data are classified into three types: protein-ligand complex structures, small-molecule structures, and protein sequences. PhysDock is evaluated using three benchmark datasets: PoseBusters, DeepDockingDare, and PhiBench. **Protein-ligand Complex Structures** Protein–ligand complex structures are derived from the Plinder dataset^24^, which includes over 400,000 protein–ligand interaction (PLI) systems from the Protein Data Bank (PDB) up to June 2024, spanning a broad range of protein families and small-molecule types. The data curation process for each PLI system involves the following steps: (i) extract structural data from PDB entries; (ii) generate biological assemblies using OpenStructure^51^; (iii) detect ligand-like chains and their interactions with proteins using PLIP^33^; and (iv) curate these interactions based on specific criteria such as chain type, and the number of residues. Following curation, each Plinder system comprises a receptor composed of one or more protein chains and multiple ligands, including small molecules, ions, peptides, and nucleic acid chains. To further enhance structural quality and prevent information leakage, additional filtering steps were applied to the Plinder dataset. These include the removal of systems with resolutions lower than 6Å, oligosaccharide systems, nucleic acid systems, systems with missing ligand atoms, multi-ligand systems exhibiting ligand clashes or overlaps, and any systems that are part of the PoseBusters^20^ and DeepDockingDare benchmark datasets.

### Small-Molecule Structures

To enhance PhysDock’s sensitivity to the geometric features of small molecules, the Plinder dataset was augmented with randomly sampled drug-like structures from the ZINC20 database^52^ during training.

### Protein Sequences

PhysDock leverages multiple sequence alignment (MSA) techniques to enhance protein feature extraction and improve training stability. The protein sequence databases used include the Big Fantastic Database^21^ (BFD), Uniclust30^53^ v.2018_08, UniRef90^54^ v.2020_01, MGnify^55^ v.2018_12, and UniProt^56^ (Swiss-Prot&TrEMBL, 2017-11). BFD and Uniclust30 were queried using HHBlits^57^ from HH-suite v.3.0-beta.3, while UniRef90, MGnify, and UniProt were searched using Jackhmmer^58^ from HMMER3^59^. The resulting MSAs were then processed into MSA features, which were used as model inputs.

### Benchmark Data

To ensure rigorous and reproducible evaluation of molecular docking methods, we established a standardized preprocessing pipeline for benchmark datasets that address common challenges such as multi-chain complexes, non-target ligands, and incomplete structural data. For each complex, the target ligand was extracted from its SDF file using explicit residue identifiers (*e*.*g*., CCD ID: LIG or UNK), while the receptor structure was refined by removing all non-protein components (*e*.*g*., ions, solvents, and redundant ligands) from the original PDB file. This process ensures consistent ligand–receptor pairing across all benchmarks. For complexes with total protein lengths exceeding 1,500 residues, we performed chain truncation to enhance computational efficiency while preserving biological relevance. Specifically, we calculated the minimum distance between each protein chain and the target ligand, retaining only chains with at least one residue within 15Å of the ligand. This strategy effectively eliminates crystallization artifacts while preserving ligand-binding interfaces. To accommodate baseline methods that require full-length protein sequences (*e*.*g*., AlphaFold3), we reconstructed missing residues using mmCIF files from the PDB database. For traditional docking tools (*e*.*g*., Vina, Glide), we provided preprocessed PDB files with explicit binding pockets to facilitate method-specific comparisons.

### Featurization

We provide a summary of the featurization strategies employed in PhysDock.

### Feature Shape Notations

PhysDock’s raw features are hierarchically organized into four sequential units: Chainwise, Conformerwise, Tokenwise, and Atomwise. Specifically, each protein chain and ligand molecule is considered as a distinct chain. For proteins, each amino acid residue acts as both a conformer and a token. For ligands, each molecule is treated as a conformer, with individual atoms serving as tokens. Features from higher-level units are inherited by all associated lower-level units. For example, each residue shares the same entity ID as its parent chain. This hierarchical design enables the propagation of high-level features to lower levels, resulting in model inputs consisting solely of Tokenwise and Atomwise features. Atomwise features for both proteins and ligands primarily originate from the geometric attributes of small molecules associated with each conformer, including atomic charge, element type, degrees of freedom, hybridization type, chirality, and the reference position of the conformer. Tokenwise features encompass covalent bond characteristics of ligands, shortest paths between atoms in the molecular graph, sequence features at the token level, MSA-derived features, C*β* distance grams, and positional enbeddings for each token. We denote the number of tokens as N_token_, the number of atoms as N_atom_, and the number of MSA rows used in the model by N_msa_.

### Target Feature and MSA Feature

The target feature (target_feat) and MSA feature (msa_feat) are token-wise features. For MSA feature, the MSA sequences are subjected to MSA Pairing. This process results in a paired MSA and Deletion Matrix, with a maximum of 16,384 rows. The one-hot encoded MSA, with dimensions [N_msa_,N_token_,32], is used to compute the MSA profile (msa_profile), which represents the distribution across residue types in the main MSA. Additionally, the mean number of deletions at each position in the MSA, denoted as deletion_mean ([N_token_]), is computed. During training and inference, 128 rows with the same row ID are randomly sampled from MSA and Deletion Matrix. The Deletion Matrix is then used to calculate two features: the has_deletion feature, a binary indicator of the presence of a deletion to the left of each position in the MSA, and the deletion_value feature, the raw deletion counts (the number of deletions to the left of each MSA position) transformed to [0, 1] using 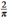 arctan 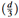. The one-hot encoded MSA, has_deletion, and deletion_value are concatenated to form the msa_feat. The target_feat is constructed by concatenating the one-hot encoded residue type, msa_profile and deletion_mean.

### Conformer Features

Each residue and ligand is associated with a corresponding small-molecule conformer, whose chemical descriptors are represented as atom-wise single features (ref_feat) and token-wise pair features (rel_tok_feat). The ref_feat encompasses 3D coordinates, atomic charge, atomic number, aromaticity, degree, hybridization type, implicit valence, chirality label, and a flag indicating whether the atom is part of a ring of a specified size. The rel_tok_feat, derived solely from ligand conformers, is based on the edge features between atom pairs within the corresponding conformer, including the shortest path length between the atom pair, whether the atoms are bonded, bond type, whether the bond is within a ring, whether the bond is conjugated, and whether the bond is directional. The node and edge features of each conformer are computed using RDKit^60^.

### Positional Features

The position-related features are token-wise single features, including residue_index, asym_id, entity_id, and sym_id. The residue_index serves as a unique identifier for each conformer. The asym_id distinguishes residues belonging to different chains. The entity_id differentiates entities with distinct sequences. The sym_id is utilized to further distinguish individual chains within the same entity. In this context, each identical ligand is treated as a separate entity and an independent chain.

### Backbone Distogram

A distogram is a binned representation of pairwise distances between atoms or residues in a protein structure, calculated using Euclidean distance and organized into bins to simplify analysis. The *β* carbons of the protein backbone (or C*α* for glycans) are used to compute the distogram, which serves as a coarse-grained structural feature of the protein. Distances ranging from 3.25 to 50.75Å are discretized into a total of 39 bins.

### Physical Priors

The physical prior information fed into the model includes the pocket residue feature (pocket_res_feat) and the key residue feature (key_res_feat), both of which are token-wise sequence features. The pocket_res_feat indicates whether a residue is part of the binding pocket. The key_res_feat identifies residues involved in six types of interactions: salt bridges, hydrogen bonds, *π*-stacking interactions, *π*-cation interactions, hydrophobic interactions, and metal complexes. In redocking, PhysDock automatically assigns residues within 10Å of any atom in the GT ligand molecule as pocket residues. During screening, residues within 10Å of any atom of a known binding small molecule in the target are considered as pocket residues. The calculation of key residues is performed using PLIP, and during inference, it is randomly activated with a 50% probability.

### Data augmentation

We summarize the data augmentation techniques applied during the training phase of PhysDock.

### Spatial Cropping

Spatial cropping is a key data augmentation technique that enhances training efficiency, reduces memory pressure, and improves training outcomes. PhysDock implements three spatial cropping methods: (i) spatial_crop randomly selects a protein residue token as the center and trims the nearest given number of tokens and atoms; (ii) the spatial_interface_crop centers on tokens within the interface, defined as tokens with other chain tokens within 15Å; (iii) spatial_ligand_crop centers on the coordinate center of the ligand. During training, PhysDock activates these three cropping methods with probabilities of 0.2, 0.4, and 0.4, respectively. Spatial_ligand_cropping can also be used to accelerate inference.

### Chirality Randomization

Each atom in the complex that possesses chirality has a 0.1 probability of being assigned an unknown chirality.

### MSA Sampling

After MSA pairing, 128 sequences are randomly selected from all MSA to form an MSA cluster.

### Center Random Augmentation

Center random augmentation is applied to the complex structure by first centering the cropped structure at 0. A uniform rotation matrix is then applied, along with a random Gaussian shift with a standard deviation of 1.

### Conformer Reference Position Randomization

The reference position features of each conformer in the structure undergo center random augmentation.

### Key Residue Feature Masking

Key residue features are activated with a 0.5 probability. When activated, each type of key interaction is randomly masked with a probability of 0.5.

### Pocket Feature Randomization

Pocket features are activated with a 0.5 probability. When activated, the determination of whether a receptor residue is a pocket residue is based on two conditions: the presence of any atom within a given cutoff range and the C*α* atom of the residue being within the cutoff. The cutoff range is randomly selected between 6 and 12Å.

### Symmetry ID Randomization

For protein chains with the same entity, the sym_id is randomly shuffled.

### Distance Gram Feature Masking

The C*β* distance gram feature of the protein backbone is activated with a 0.75 probability. When activated, the tokens of the protein chain are randomly masked with a probability ranging from 0 to 0.4 before calculating the distance gram.

### Dataset Mixing

During training, the Plinder dataset is mixed with the ZINC20 small molecule dataset, with a 0.1 probability of uniformly selecting a molecule from ZINC20^52^.

### Model architecture

The PhysDock model is composed of two major components: DiffusionConditioning and DiffusionModule. These components work synergistically to integrate multimodal features and structural priors into the diffusion-based generative process.

### DiffusionConditioning

DiffusionConditioning acts as the encoder for multimodal features and physical priors, converting atom-wise and token-wise inputs into latent representations to integrate diverse information sources. The architecture comprises two components: AtomEmbedder and TokenEmbedder. The core of AtomEmbedder is the AtomTransformer. The significant network layers of TokenEmbedder include EvolutionTransformer, TriangularTransformer, and PairTransformer. These components are designed to capture both local and global interactions within molecular structures.

### DiffusionModule

DiffusionModule takes the denoised structures from the previous step, conditioned on the latent representations encoded by DiffusionConditioning, and further refines them through a series of diffusion steps. The module consists of three diffusion transformer layers. The first and third layers are AtomDiffusionTransformer, which operate on full-atom representations to capture spatial interactions at the atomic level. The middle layer is a token-wise DiffusionTransformer, functioning similarly to a low-pass filter to integrate token-wise signals. Notably, our AtomDiffusionTransformer employs a full-atom visible attention mechanism to capture spatial interactions at the atomic level, thus enhancing the model’s ability to generate structurally accurate outputs. Notably, our AtomDiffusionTransformer employs a full-atom visible attention mechanism to capture interactions between distant atoms at the atomic level, thus enhancing the model’s ability to generate structurally accurate outputs. The DiffusionTransformer layers are designed to ensure robustness and efficiency of the diffusion process. By integrating these components, PhysDock effectively balances data-driven denoising with the enforcement of physical constraints, thereby improving the accuracy and reliability of molecular structure generation.

### Physics-guided diffusion model

#### Diffusion Models

Diffusion models are a class of generative models that iteratively refine noisy inputs to generate clean samples. The core idea is to gradually add noise to the data during the forward diffusion process and then learn to reverse this process to generate new samples. This is achieved through a reverse diffusion process, which is learned by a neural network. The reverse process can be described by a stochastic differential equation (SDE) or an ordinary differential equation (ODE), depending on the specific formulation. The forward diffusion process can be described by the following SDE:

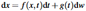

where *f* (*x, t*) is the drift coefficient, *g*(*t*) is the diffusion coefficient, and d*w* represents a standard Brownian motion. In the context of diffusion models, Song *et al*. made significant contributions to the understanding and development of these models through the lens of Stochastic Differential Equations (SDEs)^61^. Specifically, they introduced the concepts of Variance Preserving (VP) and Variance Exploding (VE) SDEs, which are two different ways to model the diffusion process. The VP-SDE is designed to maintain a constant variance throughout the diffusion process. This is achieved by carefully designing the noise schedule and the diffusion process. The VP-SDE is defined as follows:

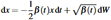

where: *x* is the state variable, *β* (*t*)is a schedule of variance parameters. d*W* is a Wiener process (standard Brownian motion). The key property of the VP-SDE is that it preserves the variance of the data distribution. Specifically, if the initial distribution *x*_0_ has unit variance, the variance of *x*_*t*_ remains constant over time. This is achieved by balancing the variance of the noise added at each step with the variance of the data. The VE-SDE allows the variance to increase exponentially over time. This leads to faster diffusion but can result in less stable training compared to the VP-SDE. The VE-SDE is defined as follows:

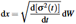

where *σ* (*t*) is an increasing function of time. The key property of the VE-SDE is that the variance of the diffusion process increases exponentially over time. This is achieved by setting the diffusion coefficient to be proportional to the derivative of the variance schedule. In non-equivariant molecular diffusion generative models, the VP-SDE (Variance Preserving Stochastic Differential Equation) framework is particularly effective due to its constant variance maintenance throughout the diffusion process. This is supported by empirical results, including those from PhysDock.

#### The EDM Framework

The EDM framework^62^, utilized in PhysDock and proposed by Karras *et al*., presents significant advancements in diffusion-based generative models. EDM proposes a unified framework to express diffusion models, enabling modular design and facilitating easier exploration of the model design space. In this framework, the data distribution of 3D coordinates of protein-ligand complex *p*_data_(*x*) is perturbed by adding Gaussian noise to create a family of distributions *p*(*x*; *σ*). The noise levels *σ* are controlled by a schedule *σ* (*t*). The probability flow ordinary differential equation (ODE) is defined as:

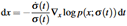

This ODE describes how the noise level evolves over time. The score function ∇_*x*_ log *p*(*x*; *σ*) is approximated by a neural network *D*_*θ*_ (*x*; *σ*), trained to minimize the expected L2 denoising error:

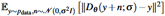

Except the the defined unified framework, EDM design karras noise schedule to optimize the sampling process by controlling the noise levels during the sampling steps. The noise levels *σ* (*t*) are chosen to be *σ* (*t*) = *t*, and the time steps *t*_*i*_ are defined using a polynomial schedule:

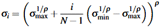

where *ρ* controls the shape of the schedule. The stochastic sampler combines deterministic steps with Langevin-like noise injection and removal. The key steps include (i) Adding noise to the sample:

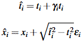

and (ii) Evaluating the score function and updating the sample:

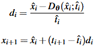

Where *γ*_*i*_ represents a scaling factor or a step size in the context of the noise schedule. *ε*_*i*_ represents a random noise vector drawn from a standard normal distribution. In PhysDock, we set *ρ* = 1000, which enhances sampling resolution at low noise levels and enables more accurate results with significantly fewer diffusion steps.

#### Physics-Guided Sampling with Iterative Projection

PhysDock integrates physics-based constraints into the diffusion process to ensure the generated samples are physically plausible. This is achieved through a combination of stepwise denoising, physics-based projection, and iterative refinement: (i) stepwise Denoising, at each reverse diffusion step t, a neural network predicts the denoised structure 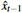 based on the current noisy input *x*_*t*_; (ii) physics-based projection, subset of denoised samples undergoes physics-based projection, where structure matching or energy minimization using a molecular mechanics force field corrects bond length and angle violations, steric clashes, and other inconsistencies, yielding a physically plausible structure 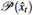; (iii) iterative refinement, the corrected conformation 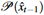 is fed back into the diffusion process as a conditioning for subsequent steps, ensuring consistency with both the learned data distribution and physical priors.

#### Projection Schemes

The exploring-matching-aligning-replacing strategy is used for ligand projection under high noise conditions. This strategy involves: (i) exploring a wide range of ligand conformations to comprehensively cover the conformational space *via* sampling; (ii) matching the denoised ligand with the ligands in the sampled conformer library; (iii) aligning the closest conformer onto the denoised ligand structure; (iv) replacing the denoised ligand structure with the aligned conformer. This strategy balances high noise levels with the stereochemical constraints of the ligand molecules. The relaxing-aligning-replacing strategy is used for ligand projection under low noise conditions. This strategy involves: (i) relaxing to optimize the denoised structure using MMFF; (ii) aligning the optimized structure to the target; (iii) replacing the denoised molecule structure with the aligned and optimized one. This strategy ensures that the generated structures are optimized based on MMFF under low noise conditions. Both conformer generation and force field optimization are performed using the RDKit toolkit^60^.

#### Adaptive Projection Scheduling

Physical corrections are selectively applied during the reverse diffusion process. Early steps with high noise (t ≥*η*) prioritize data-driven denoising to capture global structural patterns, while later steps (t<100) focus on local refinements guided by physical principles. This scheduling avoids over-constraining the model during high-noise phases, where premature corrections could distort the generative distribution.

### Clustering and ranking

We performed K-means clustering and ranking on the sample sets obtained from multiple rounds of sampling within the same system. The clustering process involved two main steps: similarity calculation and cluster partitioning. First, we employed PAL-RMSD as the similarity metric between samples, then partitioned the sample set into a predefined number of clusters, *K*. For each cluster, we selected the nearest neighbor to the cluster center as the representative sample. These representative samples were ranked based on their distance to the global centroid of all samples. In the three benchmark tests of PhysDock, we selected five representative samples from the sorted clusters, *i*.*e*., *K* = 5.

## Extended data

**Extended Data Fig. 1.**
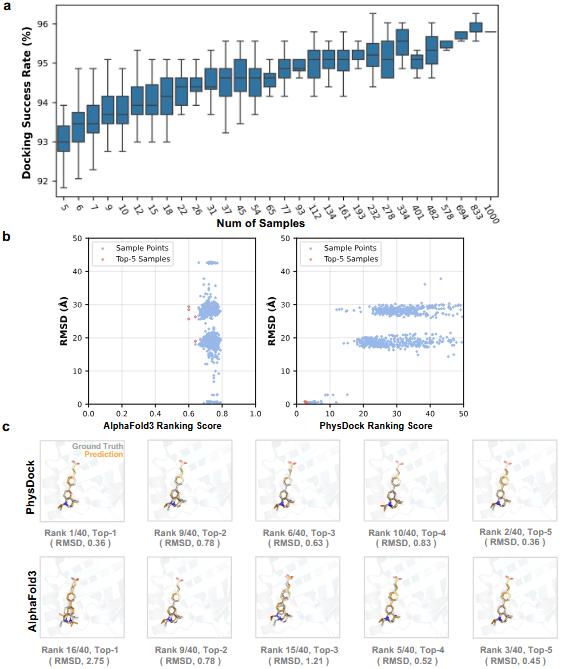
Comparison of ranking methods between AlphaFold3 and PhysDock using AlphaFold3-generated predictions on the PoseBusters dataset. **(a)** Top 5 docking success rate (PAL-RMSD<2Å) of PhysDock as a function of the number of generated samples. **(b)** Left: AlphaFold3’s ranking score vs. PAL-RMSD plot; Right: PhysDock’s ranking score vs. PAL-RMSD plot. Each point represents one of the 1000 samples generated by AlphaFold3 (PDB ID: 7AA0, using seed 41 and 42), where the x-axis shows the ranking score (lower is better) and the y-axis indicates RMSD to the ground truth. Red points denote the Top-5 ranked samples. **(c)** Top 5 AlphaFold3 predictions for PDB ID: 7AA0 (from 40 samples generated using seed 42) ranked using PhysDock scores and a modified AlphaFold3 ranking score(1.1 − AlphaFold3 ranking score).

**Extended Data Fig. 2.**
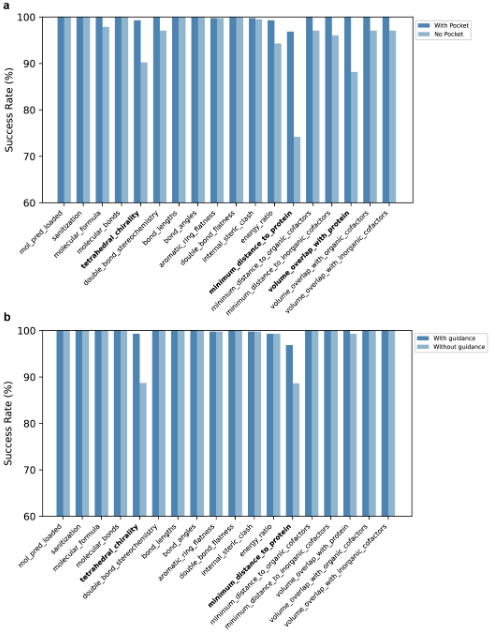
Success rates of PhysDock on the PoseBusters dataset evaluated across 18 PB validation criteria. Performance is shown for conditions with and without prior pocket information, and with and without two-phase physics guidance. Validation checks with lower pass rates are highlighted in bold.

**Extended Data Fig. 3.**
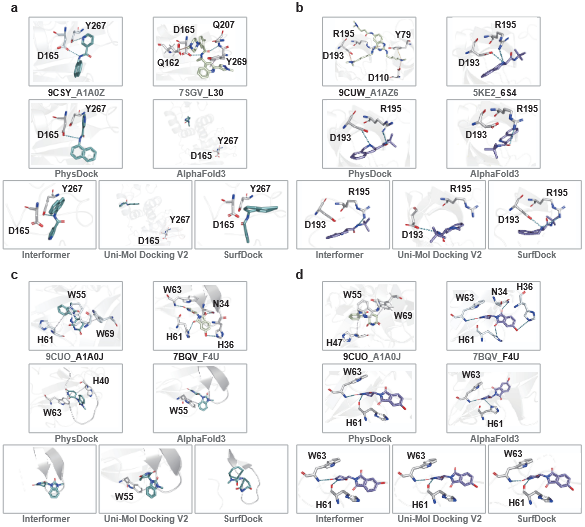
Representative cross-docking cases comparing the predicted binding modes from various models, including PhysDock, AlphaFold3, Interformer, Uni-Mol Docking V2, and SurfDock, with the ground truth. Each case involves docking a ligand from one protein structure into a different protein structure that was resolved in complex with another ligand.

**Extended Data Fig. 4.**
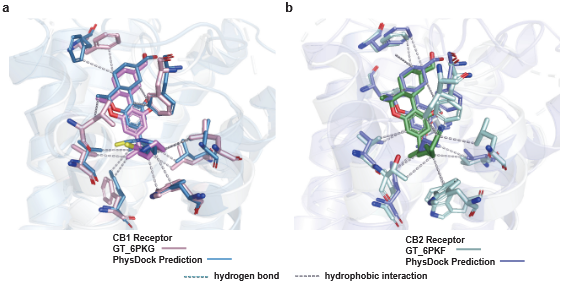
Overlay of binding modes predicted by PhysDock (blue for CB1, purple for CB2) and the ground truth (pink for CB1, green for CB2). Dashed lines represent hydrogen bonds (cyan) and hydrophobic interactions (gray).

**Extended Data Fig. 5.**
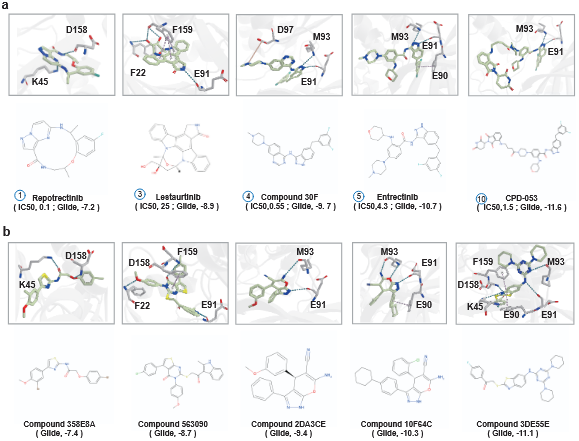
**(a)** Docking poses of known NTRK3 kinase inhibitors. This panel supplements Figure 6d by presenting additional inhibitors not included in the main figure. Corresponding 2D structures are displayed below. **(b)** Docking poses of high-affinity SPECS hits for NTRK3 kinase. Corresponding 2D structures are displayed below. Hydrophobic interactions are omitted for clarity.

**Extended Data Fig. 6.**
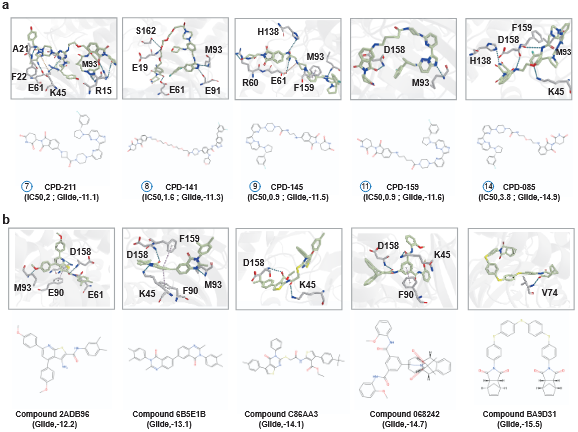
**(a)** Docking poses of known NTRK3 kinase inhibitors. This panel supplements Figure 6d by presenting additional inhibitors not included in the main figure. Corresponding 2D structures are displayed below. **(b)** Docking poses of high-affinity SPECS hits for NTRK3 kinase. Corresponding 2D structures are displayed below. Hydrophobic interactions are omitted for clarity.

**Extended Data Table 1.** The compile-free inference times of PhysDock and Chai-1, and the compiled inference time of AlphaFold3, represent the average time required to generate 40 samples per method, averaged across multiple independent runs. As AlphaFold3 is JAX-based and requires compilation prior to inference, its reported timing reflects only the compiled inference speed, with compilation overhead excluded.

| Methods | 1 H20 96GB (GPU secs)<sup>1</sup> |  |  |
| --- | --- | --- | --- |
|  | 384 Tokens | 768 Tokens | 1024 Tokens |
| Chai-1 | 296 | 644 | 913 |
| AlphaFold3 | 75 | 123 | 187 |
| PhysDock | 18 | 67 | 102 |

## Data availability

The protein sequence data used in our study are all sourced from public datasets. The curated training, validation, and benchmark datasets are available *via* Zenodo at https://zenodo.org/records/15178859, https://zenodo.org/records/15206943, https://zenodo.org/records/15209515 and https://zenodo.org/records/15220255.

## Code availability

The source code used to run inference and generate the results shown in this study is available under an MIT License via GitHub at https://github.com/KexinZhangResearch/PhysDock.

## Acknowledgments

We appreciate the discussions with Suwen Zhao and Jianjun Cheng from ShanghaiTech University, as well as Zhenting Gao and Xianqiang Song from Structure Therapeutics. We gratefully acknowledge the support from Shanghai Frontiers Science Center for Biomacromolecules and Precision Medicine. We also thank the High-Performance Computing (HPC) platform at ShanghaiTech University for providing computational resources, and Cellverse Co., Ltd. for their technical support.

## Notes

https://github.com/KexinZhangResearch/PhysDock

https://zenodo.org/records/15178859

https://zenodo.org/records/15206943

https://zenodo.org/records/15209515

https://zenodo.org/records/15220255

### Competing Interest Statement

The authors have declared no competing interest.

### Summary of Updates

Author affiliations updated; Supplemental files updated.

