## Supplementary Information for "PhysDock: A Physics-Guided All-Atom Diffusion Model for Protein-Ligand Complex Prediction"

**3 Loss** **12**
**4 Training details** **12**
**5 Baseline details** **13**
**6 Supplementary Figures** **14**
**7 Supplementary Tables** **20**
**References** **37**

### 1 Additional data details

Featurization and data augmentation are described in the Methods section of the main text. The MSA featurization pipeline is adapted from AlphaFold2, while the MSA pairing scheme follows that of AlphaFold3. Additional details on data and feature processing are provided below.

#### 1.1 Filtering

We extensively processed the Plinder dataset to construct the training and validation sets for PhysDock, as follows:

1. Removing systems with a resolution greater than 9 Å.

2. Removing systems containing RNA, DNA, and peptides.

3. Removing systems with ligands that have missing atoms.

4. Removing systems present in the benchmark dataset.

5. Removing systems with only a ligand or only a receptor.

6. Removing systems in the bio assembly where atoms, especially ligand atoms, have clashes (non-hydrogen atom distances less than 1 Å).

Ultimately, we obtained approximately 260,000 training and validation systems after filtering.

#### 1.2 Metadata Processing

To accelerate dataset loading, we saved the following metadata:

1. `pdb_id_meta_data`: storing the structure determination method (*e.g.*, x-ray diffraction), release date, resolution, and all PDB chain IDs for each system.

2. `chain_id_meta_data`: storing the sequence of the specified chain_id, the sequence3 composed of CCD identifiers, and the MD5 code of the sequence. The MD5 code is used to retrieve existing MSA features and uniprot MSA features.

3. `ccd_id_meta_data`: storing all conformer features corresponding to each CCD conformer.

4. `train_val_sample_id_meta_data`: storing the sampling weight assigned to each training sample.

The weight for each sample is calculated based on the combined weight of the receptor and ligand. The receptor weight, ($W_{\text{receptor\_identity}}$), is determined by sequence identity, using a 40% sequence identity threshold. The ligand weight, ($W_{\text{ligand\_identity}}$) uses a 100% identity (*i.e.*, the weight is identical for ligands that are exactly the same).

$$W = W_{\text{receptor}} \times W_{\text{ligand}} = W_{\text{receptor\_identity}} \times W_{\text{ligand\_identity}}$$

The resulting weights are then used to guide the selection of the entire PDB structure for training, as well as to determine the receptor and ligand for sample localization. Additionally, they are used to process the model input for training according to the cropping strategy.

### 2 Model details

The network architecture and pseudocode (shown below) incorporate key innovations from AlphaFold2, AlphaFold3, Llama3, and Stable Diffusion 3, but is specifically tailored for molecular docking tasks.

#### 2.1 Model Inputs

The input features of the model can be broadly categorized into sequence-related features, conformer-related features, backbone-related features, physical priors, and positional features. Below, we provide a brief description of each of these main feature categories.

| Input Feature | Shape | Feature Type | Description |
|---|---|---|---|
| msa_feat | $[N_{msa}, N_{token}, 32]$ | Sequence Feature | MSA-related features obtained from the processing of MSA and deletion matrix. |
| target_feat | $[N_{msa}, N_{token}, 32]$ | Sequence Feature | Sequence-related features derived from the processing of the sequence and MSA profile. |
| ref_feat | $[N_{atom}, 32]$ | Conformer Feature | Reference conformer node-related features generated for each residue or ligand by RDKit. |
| ref_pos | $[N_{atom}, 3]$ | Conformer Feature | Randomly generated coordinated by RDKit. |
| ref_space_uid | $[N_{atom}]$ | Conformer Feature | A unique ID of each conformer to distinguish atoms in different conformers. |
| rel_tok_feat | $[N_{token}, N_{token}, 32]$ | Conformer Feature | Edge-related features for the reference conformer of ligand, zero for the receptor. |
| token_bonds | $[N_{msa}, N_{token}, 32]$ | Conformer Feature | Covalent bonds for each atom in the reference conformer. |
| dgram_feat | $[N_{token}, N_{token}, 40]$ | Backbone Feature | Distance Gram features computed for backbone C$\beta$ atoms. |
| key_res_feat | $[N_{token}, 7]$ | Physical Priors | Indicators to mark whether a token is a key residue. |
| pocket_res_feat | $[N_{token}]$ | Physical Priors | Indicators to mark whether a token is a pocket residue. |
| residue_index | $[N_{token}]$ | Position Feature | Residue index in each chain. |
| asym_id | $[N_{token}]$ | Position Feature | Unique chain IDs. |
| entity_id | $[N_{token}]$ | Position Feature | Unique entity IDs. |
| sym_id | $[N_{token}]$ | Position Feature | IDs to distinguish different chains of the same entity. |

**Table 1.** Details of model input features.

#### 2.2 Primitives

Below, we will present the primary network layers that constitute PhysDock. All attention mechanisms incorporate a gating mechanism, and some attention mechanisms introduce QK norm to stabilize the numerical values of the latent representations.

**Algorithm 1** AdaLayerNormZero

**def** AdaLayerNormZero($\{x_i\}\{t\}$)

$t \leftarrow \mathrm{silu}(t)$

$x_i \leftarrow \mathrm{LayerNorm}(s, \mathrm{affine} = \mathrm{False})$

$shift \leftarrow \mathrm{Linear}(t)$

$scale \leftarrow \mathrm{Linear}(t)$

$gate \leftarrow \mathrm{Linear}(t)$

$x_i \leftarrow x_i \odot (1 + scale) + shift$

**return** $\{x_i\}, \{gate\}$

**Algorithm 2** FeedForward

**def** FeedForward($\{x_i\}$)

$x \leftarrow \mathrm{silu}(\mathrm{Linear}(x)) \odot \mathrm{Linear}(x)$

$x \leftarrow \mathrm{Linear}(x)$

**return** $\{x_i\}$

**Algorithm 3** Transition

**def** Transition($\{x_i\}$)

$x_i \leftarrow \mathrm{RMSNorm}(x_i)$

$x_i \leftarrow \mathrm{FeedForward}(x_i)$

**return** $\{x_i\}$

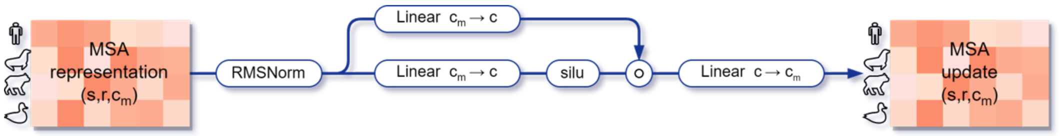

**Figure 1.** MSATransition layer.

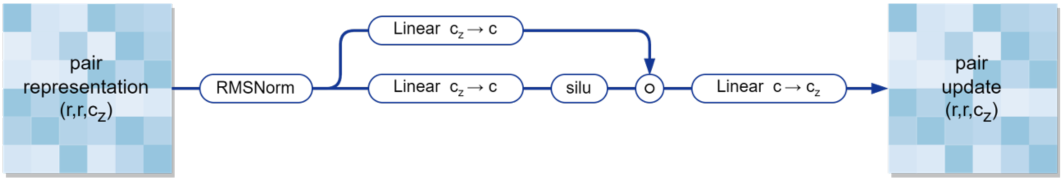

**Figure 2.** PairTransition layer.

---

**Algorithm 4** DiTTransition

---

**def** DiTTransition($\{x_i\}, \{t\}$)

$x_i, g_i \leftarrow$ AdaLayerNormZero$(x_i, t)$

$x_i \leftarrow$ FeedForward$(x_i) \odot g_i$

**return** $\{x_i\}$

---

**Algorithm 5** ScaledDotProductAttention

---

**def** SDPA($\{x_i\}\{b_{ij}^h\}$)

$q_i^h \leftarrow \mathrm{Linear}(x)$

$k_i^h \leftarrow \mathrm{Linear}(x)$

$v_i^h \leftarrow \mathrm{Linear}(x)$

$a_{ij}^h \leftarrow \mathrm{softmax}_j \left( \frac{1}{\sqrt{c}} q_i^{h\top} k_j^h + b_{ij}^h \right)$

$x_i \leftarrow \sum_j a_{ij}^h c_j^h$

**return** $\{x_i\}$

---

**Algorithm 6** ScaledDotProductAttentionNobias

---

**def** SDPANoBias($\{x_i\}$)

$q_i^h \leftarrow \mathrm{Linear}(x)$

$k_i^h \leftarrow \mathrm{Linear}(x)$

$v_i^h \leftarrow \mathrm{Linear}(x)$

$a_{ij}^h \leftarrow \mathrm{softmax}_j \left( \frac{1}{\sqrt{c}} q_i^{h\top} k_j^h \right)$

$x_i \leftarrow \sum_j a_{ij}^h c_j^h$

**return** $\{x_i\}$

---

**Algorithm 7** AttentionWithPairBias

---

**def** AttentionWithPairBias($\{x_i\}, \{t\}$)

$x_i \leftarrow \mathrm{RMSNorm}(x_i)$

$b_{ij}^h \leftarrow \mathrm{RMSNorm}(z_{ij})$

$g_i \leftarrow \mathrm{Linear}(x_i)$

$x_i \leftarrow$ SDPA$(x_i, z_{ij}^h) \odot g_i$

**return** $\{x_i\}$

---

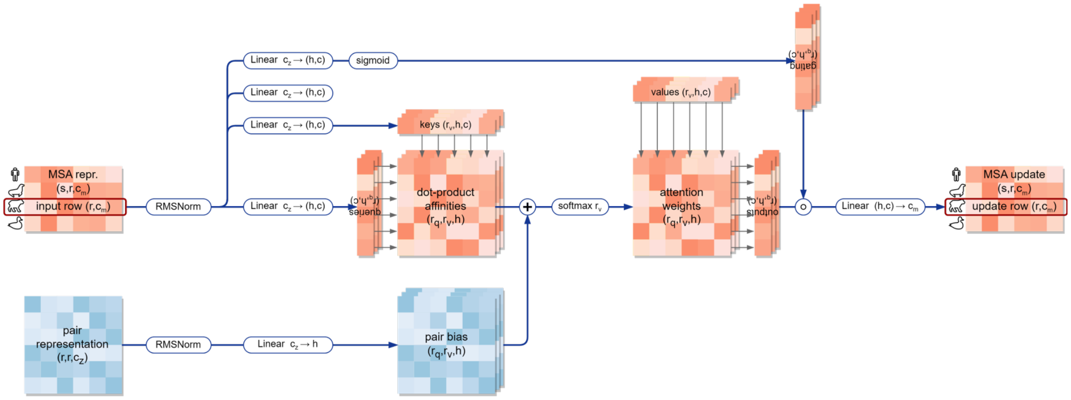

**Figure 3.** MSARowAttentionWithPairBias.

**Algorithm 8** MSAColumnAttention

**def** MSAColumnAttention($\{m_{si}\}$)

$m_{si} \leftarrow \mathrm{RMSNorm}(m_{si})$
$m_{si} \leftarrow \mathrm{SPDANobias}({m_{si}}^{\top})^{\top}$
**return** $\{m_{si}\}$

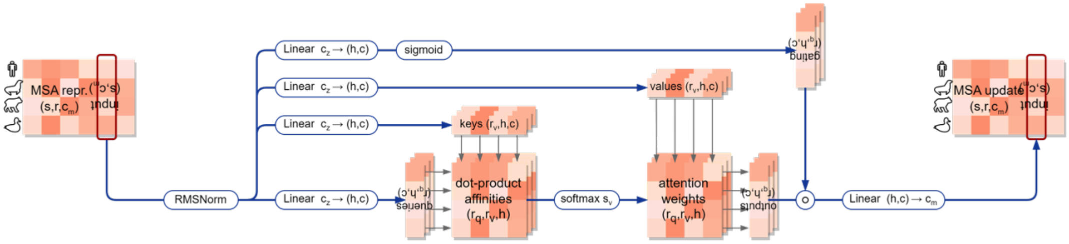

**Figure 4.** MSA Column attention.

**Algorithm 9** TriangleAttention

**def** TriangleAttention($\{z_{ij}\}$, transpose = False)

**if** transpose **then**
  $z_{ij} \leftarrow z_{ji}$
**end if**
$z_{ij} \leftarrow \mathrm{RMSNorm}(z_{ij})$
$b_{ij}^{h} \leftarrow \mathrm{Linear}(z_{ij})$
$g_{ij} \leftarrow \mathrm{Linear}(z_{ij})$
$z_{ij} \leftarrow \mathrm{SDPA}(z_{ij}, b_{ij}^{h})$
$z_{i}j \leftarrow z_{ij} \odot g_{ij}$
**if** transpose **then**
  $z_{ij} \leftarrow z_{ji}$
**end if**
**return** $\{z_{ij}\}$

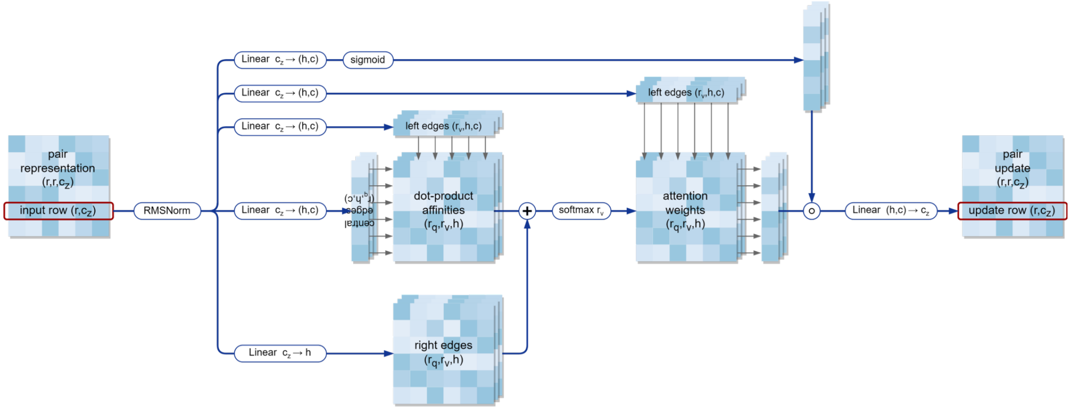

**Figure 5.** TriangleAttention layer.

---

**Algorithm 10** DiTAttention

**def** DiTAttention($\{x_i\},\{z_{ij}\},\{t\}$)

$x_i, g_i \leftarrow$ AdaLayerNormZero$(x_i, t)$

$b^h_{ij} \leftarrow \text{Linear}(\text{LayerNorm}(z_{ij}))$

$x_i \leftarrow$ SDPA$(x_i, b^h_{ij})$

$x_i \leftarrow x_j \odot g_i$

**return** $\{x_i\}$

---

**Algorithm 11** TriangleUpdate

**def** TriangleUpdate($\{z_{ij}\}$, transpose = False)

**if** transpose **then**

    $z_{ij} \leftarrow z_{ji}$

**end if**

$z_{ij} \leftarrow \text{RMSNorm}(z_{ij})$

$q_{ij} \leftarrow \text{Linear}(z_{ij}) \odot \text{sigmoid}(\text{Linear}(z_{ij}))$

$k_{ij} \leftarrow \text{Linear}(z_{ij}) \odot \text{sigmoid}(\text{Linear}(z_{ij}))$

$g_{ij} \leftarrow \text{sigmoid}(\text{Linear}(z_{ij}))$

$z_{iI} \leftarrow \sum_j z_{ij} z_{Ij}$

$z_{ij} \leftarrow \text{Linear}(\text{RMSNorm}(z_{ij})) \odot g_{ij}$

**if** transpose **then**

    $z_{ij} \leftarrow z_{ji}$

**end if**

**return** $\{z_{ij}\}$

---

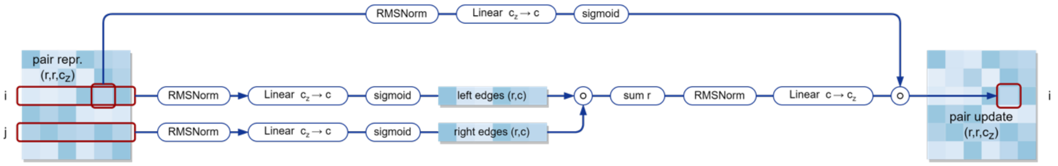

**Figure 6.** TriangleUpdate layer.

**Algorithm 12** OuterProductMean

**def** OuterProductMean($\{m_{si}\}$)

$m_{si} \leftarrow \text{RMSNorm}(m_{si})$

$q_{si} \leftarrow \text{Linear}(m_{si})$

$k_{si} \leftarrow \text{Linear}(m_{si})$

$z_{ij} \leftarrow \sum_{s} q_{si} k_{sj}$

$z_{ij} \leftarrow \text{RMSNorm}(z_{ij})$

**return** $\{z_{ij}\}$

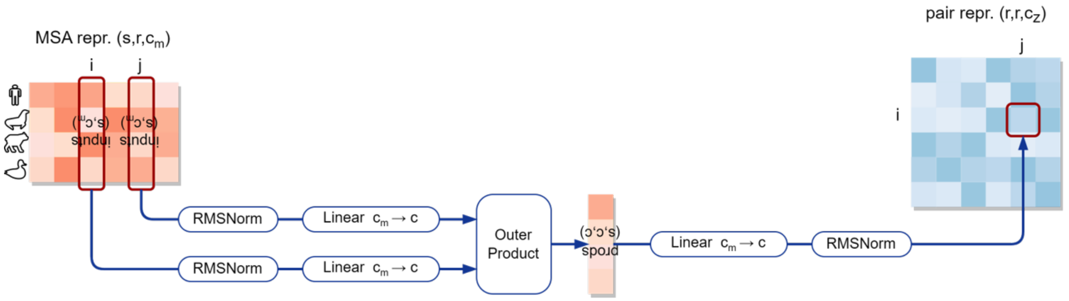

**Figure 7.** OuterpRroductMean layer. Instead of directly calculating the mean, normalization is employed in the final step.

**Algorithm 13** Timesteps

**def** Timesteps($\{t\}$, $num\_channels$, $flip\_sin\_to\_cos$, $downscale\_freq\_shift$, $scale$)

$half\_dim \leftarrow num\_channels // 2$

$exponent \leftarrow -\log(\text{max\_period}) \cdot \text{arange}(0, half\_dim) / (half\_dim - downscale\_freq\_shift)$

$emb\_base \leftarrow \exp(exponent)$

$emb \leftarrow t[:, \text{None}] \cdot emb\_base[\text{None}, :]$

$emb \leftarrow scale \cdot emb$

$emb \leftarrow \text{concat}(\sin(emb), \cos(emb), \text{dim} = -1)$

**if** flip_sin_to_cos **then**

  $emb \leftarrow \text{concat}(emb[:, half\_dim :], emb[:, : half\_dim], \text{dim} = -1)$

**end if**

**if** num_channels % 2 == 1 **then**

  $emb \leftarrow \text{pad}(emb, (0, 1, 0, 0))$

**end if**

**return** $\{emb\}$

**Algorithm 14** TimestepEmbeddings

**def** TimestepEmbeddings($\{t\}$)

$timesteps\_proj \leftarrow$ Timesteps$(t)$

$conditioning \leftarrow \text{Linear}(\text{SiLU}(timesteps\_proj))$

**return** $\{conditioning\}$

#### 2.3 Layers

Below, we will illustrate the connection methods of all the key network layers.

**Algorithm 15** AtomTransformer

```
def AtomTransformer({s_i}, {z_ij}, N_blocks)
for all l ∈ [1, ..., N_blocks] do
    s_i += AttentionWithPairBias(s_i, z_ij)
    s_i += Transition(s_i)
end for
return {s_i}
```

**Algorithm 16** RelPosEmbedder

```
def RelPosEmbedder({asym_id}, {sym_id}, {entity_id}, {residue_index}, {rel_tok_feat})
s_max ← 2                                                    # Window of chain index
r_max ← 32                                                   # Window of residue index
chain_same ← (asym_id[..., None] == asym_id[..., None, :])
entity_same ← (entity_id[..., None] == entity_id[..., None, :])
residue_offset ← residue_index[..., None] − residue_index[..., None, :] + r_max
clipped_residue_offset ← clamp(residue_offset, min = 0, max = 2 · r_max)
d_res ← where(chain_same, clipped_residue_offset, 2 · r_max + 1)
rel_pos_feat ← one_hot(d_res, arange(0, 2 · r_max + 2))
chain_offset ← sym_id[..., None] − sym_id[..., None, :] + s_max
clipped_chain_offset ← clamp(chain_offset, 0, 2 · s_max)
d_chain ← where(or(chain_same, not(entity_same)), 2 · s_max + 1, clipped_chain_offset)
rel_chain_feat ← one_hot(d_chain, arange(0, 2 · s_max + 2))
rel_feat ← concat(rel_pos_feat, rel_tok_feat, entity_same[..., None], rel_chain_feat)
return Linear(rel_feat)
```

**Algorithm 17** EvolutionTransformer

```
def EvolutionTransformer({m_si}, {z_ij}, N_blocks)
for all l ∈ [1, ..., N_blocks] do
    m_si += AttentionWithPairBias(m_si, z_ij)
    m_si += MSAColumnAttention(m_si)
    m_si += Trainsition(m_si)
    z_ij += OuterProductMean(m_si)
    z_ij += TriangleUpdate(z_ij, transpose = False)
    z_ij += TriangleUpdate(z_ij, transpose = True)
    z_ij += TriangleAttention(z_ij, transpose = False)
    z_ij += TriangleAttention(z_ij, transpose = True)
    z_ij += Trainsition(z_ij)
end for
return {m_si}{z_ij}
```

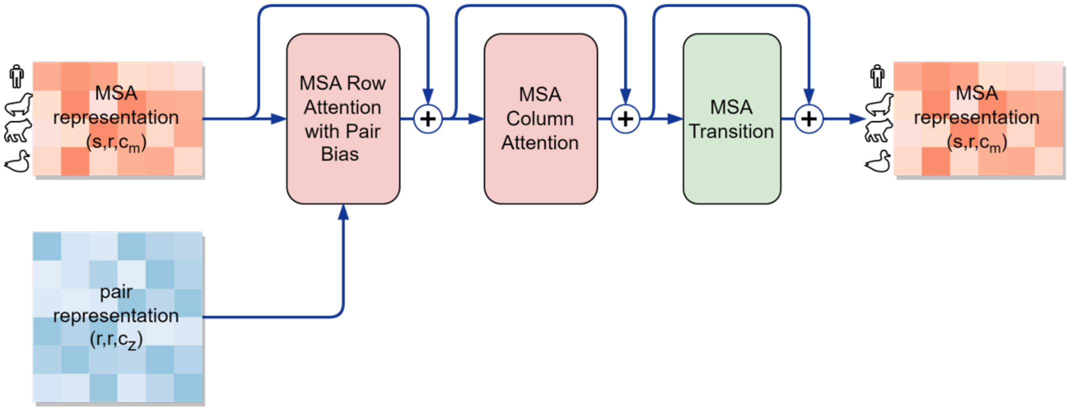

**Figure 8.** MSABlock inside EvolutionTransformerBlock.

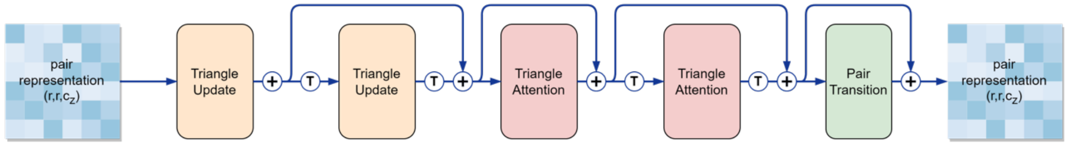

**Figure 9.** TriangleTransformerBlock.

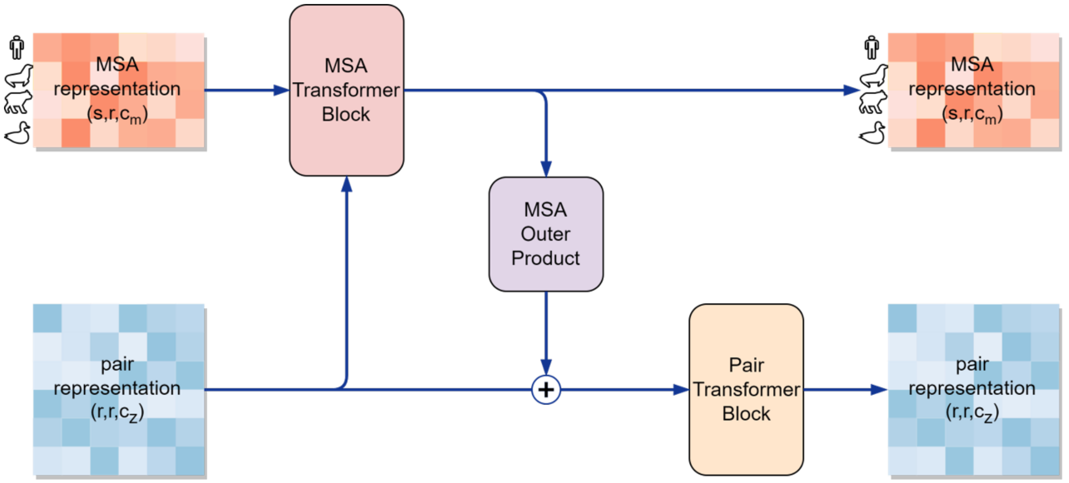

**Figure 10.** Evolution Transformer Block.

---

**Algorithm 18** TriangleTransformer

---

**def** TriangleTransformer($\{z_{ij}\}, N_{\text{blocks}}$)

**for all** $l \in [1, \ldots, N_{\text{blocks}}]$ **do**

  $z_{ij}$ += TriangleUpdate($z_{ij}$, transpose = False)

  $z_{ij}$ += TriangleUpdate($z_{ij}$, transpose = True)

  $z_{ij}$ += TriangleAttention($z_{ij}$, transpose = False)

  $z_{ij}$ += TriangleAttention($z_{ij}$, transpose = True)

  $z_{ij}$ += Trainsition($z_{ij}$)

**end for**

**return** $\{z_{ij}\}$

---

**Algorithm 19** PairTransformer

**def** PairTransformer($\{s_i\}, \{z_{ij}\}, N_{\text{blocks}}$)

**for all** $l \in [1, \ldots, N_{\text{blocks}}]$ **do**
  $z_{ij}$ += TriangleUpdate($z_{ij}$, transpose = False)
  $z_{ij}$ += TriangleUpdate($z_{ij}$, transpose = True)
  $z_{ij}$ += TriangleAttention($z_{ij}$, transpose = False)
  $z_{ij}$ += TriangleAttention($z_{ij}$, transpose = True)
  $z_{ij}$ += Trainsition($z_{ij}$)
  $s_i$ += AttentionWithPairBias($s_i, z_{ij}$)
  $s_i$ += Transition($s_i$)
**end for**
**return** $\{s_i\}\{z_{ij}\}$

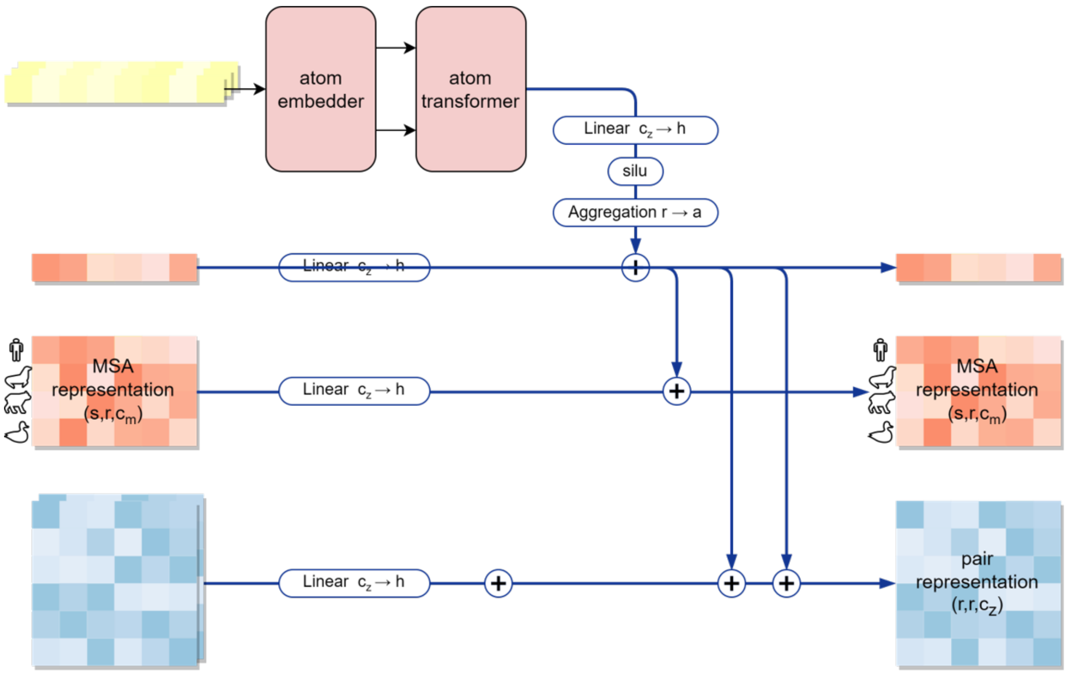

**Figure 11.** Update Atomwise representations after PairTransformer.

**Algorithm 20** DiffusionTransformer

**def** DiffusionTransformer($\{s_i\}, \{z_{ij}\}, \{t\}, N_{\text{blocks}}$)

**for all** $l \in [1, \ldots, N_{\text{blocks}}]$ **do**
  $s_i$ += DiTAttention($s_i, z_{ij}, t$)
  $s_i$ += DitTransition($s_i, t$)
**end for**
**return** $\{s_i\}$

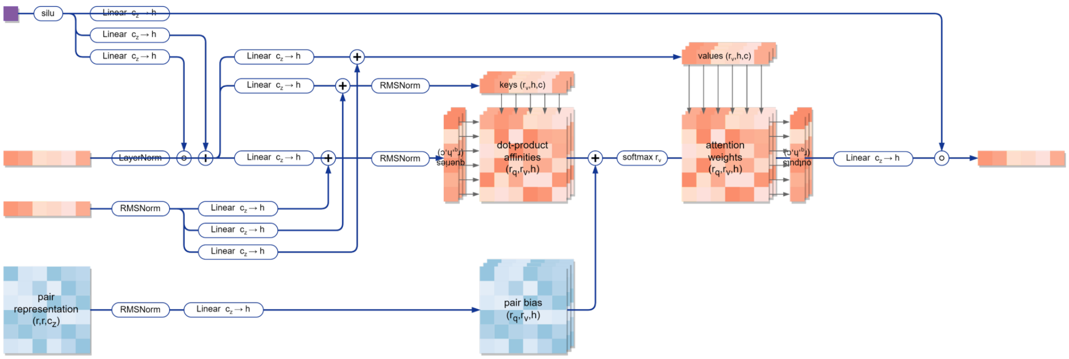

**Figure 12.** Tokenwise Diffusion Transformer Attention.

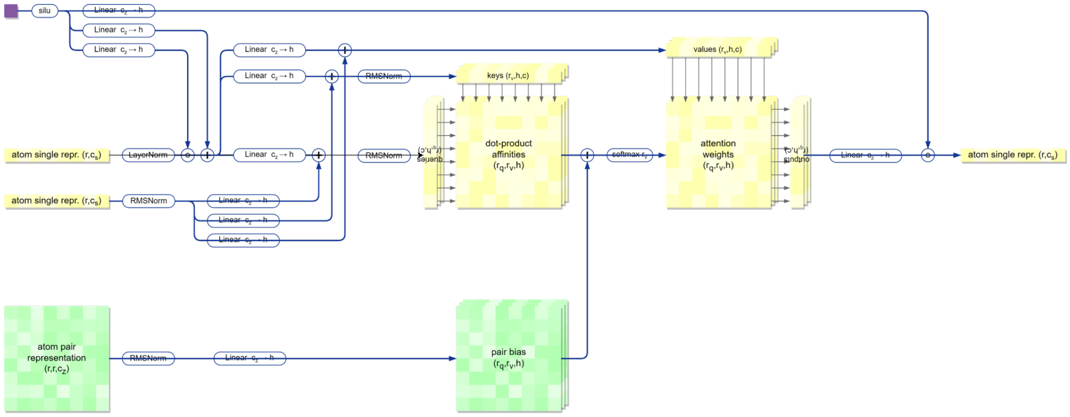

**Figure 13.** Atomwise Diffusion Transformer Attention.

**Algorithm 21** TemplatePairEmbedder

**def** TemplatePairEmbedder($\{z\}$, dgram_feat)

$z \leftarrow \text{Linear}(\text{RMSNorm}(z)) + \text{Linear}(\text{dgram\_feat})$

$z \leftarrow$ TriangleTransformer$(z)$

$z \leftarrow \text{Linear}(\text{ReLU}(\text{RMSNorm}(z)))$

**return** $\{z\}$

#### 3 Loss

Our model employs only the diffusion loss as the final loss function $\mathscr{L}_{\text{loss}}$, which integrates the Mean Squared Error (MSE) and the smooth LDDT loss:

$$\mathscr{L}_{\text{loss}} = \alpha_{\text{diffusion}} \cdot (\hat{t}^2 + \sigma_{\text{data}}^2)/(\hat{t} \cdot \sigma_{\text{data}})^2 \cdot \mathscr{L}_{\text{MSE}} + \alpha_{\text{diffusion}} \cdot \mathscr{L}_{\text{smooth\_lddt}}$$

where $\alpha_{\text{diffusion}}$ is a constant equals 4.0, $\sigma_{\text{data}}$ is a constant equals 16.0.

#### 4 Training details

Training was conducted in two phases with a maximum crop size of 384: an initial 50,000 steps using bfloat16 (bf16) mixed precision with a batch size of 48, followed by 200,000 steps using full-precision (fp32) training with the same batch size.

#### 5 Baseline details

In this study, baseline predictions were generated using existing docking methods, each producing 40 binding modes. For classical docking tools, AutoDock Vina v1.2.3 was executed with an exhaustiveness of 32, a ligand binding site threshold of $20 \sim 25$Å a protein–ligand threshold of 4Å a docking box size of 25Å× 25Å × 25Å and a grid spacing of 1ÅĠlide was performed as follows: (i) protein structures were optimized using the Protein Preparation in the Maestro module of Schrödinger (v2022-3), including hydrogen addition, bond order assignment, completion of missing side chains and loops, removal of water molecules located more than 5 Å away from the ligand, optimization of the hydrogen-bond network, and energy minimization using the OPLS-2005 force field until the heavy-atom RMSD converged to 0.3 Å ; (ii) ligand preparation was carried out using LigPrep, ultilizing the Epik module to generate possible protonation and ionization states at a target pH of $7.0 \pm 2.0$; (iii) conformers were generated, and the one with the lowest energy—based on the OPLS-2005 force field—was selected as the input for subsequent docking experiments; (iv) receptor grids were constructed using the Receptor Grid Generation module in Schrödinger, with an inner box (centered on the co-crystallized ligand) of 10 Å × 10 Å × 10 Å and an outer box extending 10Å in all directions beyond the inner box; and (v) ligand–protein docking was performed using the Glide module in standard precision (SP) mode.

For semi-flexible DL models, SurfDock first identifies amino acid residues within an 8Å radius of the target ligand to define the binding surface, computes physical features (including molecular surface, charge, hydrophobicity, and curvature), and generates a PLY file for subsequent sampling. Interformer produces an initial UFF-optimized ligand conformer, and independently defines the docking pocket as residues within a 10Å distance from the target ligand. Uni-Mol Docking V2 (Uni-Mol version 1.0.0) takes both the ligand and the full complex structure as input, selects a region within a 10Å radius of the target ligand to define the docking center and grid size, and utilizes the UFF-optimized ligand from Interformer.

For flexible DL models, NeuralPLexer operates with default settings—batched structure sampling (40 samples, 40 steps, block size 10) combined with Langevin simulated annealing. AlphaFold v3.0.0 was processed with seed 42 to generate 40 structures per complex and Chai-1 v0.5.2 was used in the MSA input mode without ESM representations. Both flexible DL models use protein sequence and ligand SMILES notation to predict the protein–ligand complex. All experiments were performed under a uniform setup, with every method executed using its default parameters to ensure consistency.

#### 6 Supplementary Figures

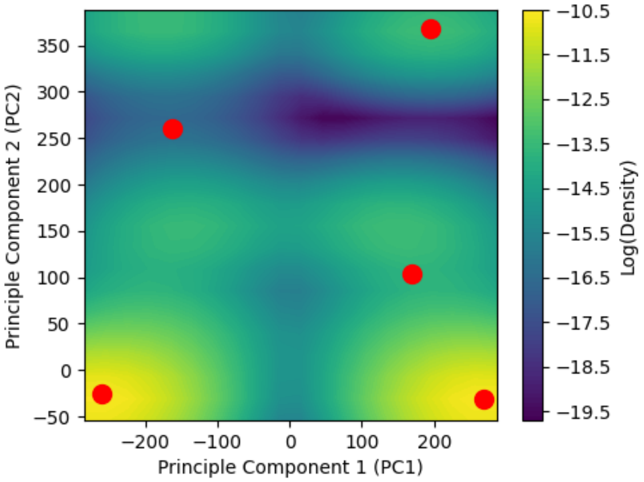

**Figure 14.** Log-density visualization of 1,000 AlphaFold3-generated samples (PDB ID: 7AA0, seeds 41 and 42). The x- and y-axes represent the first and second principal components of the RMSD distance matrix, respectively. Red points denote the Top-5 ranked samples.

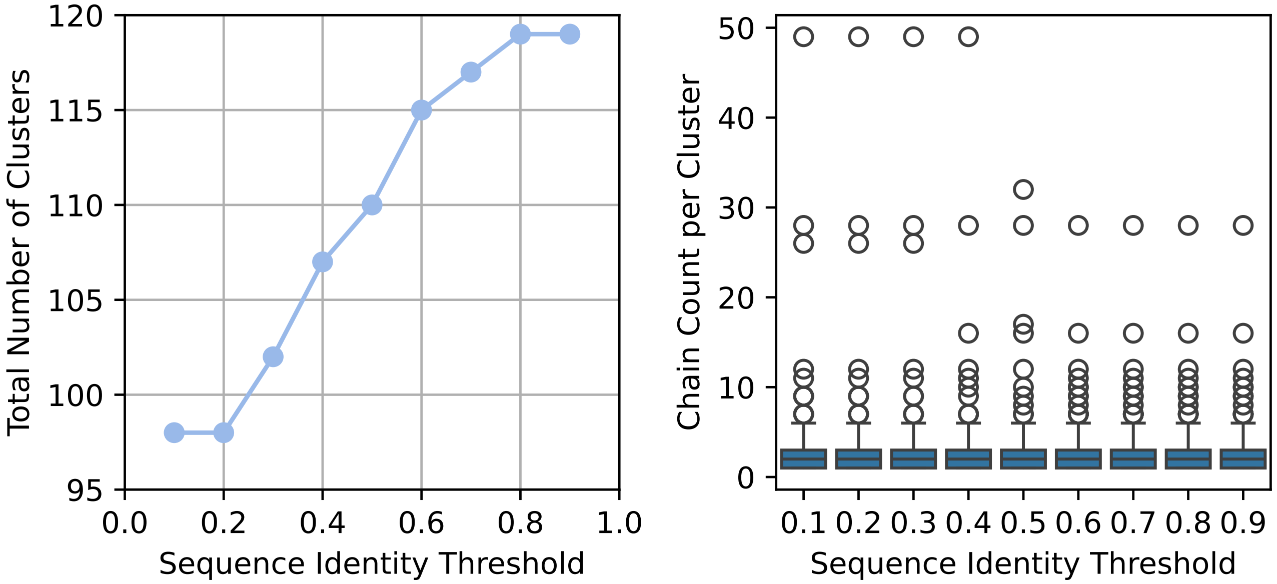

**Figure 15.** Analysis of the PhiBench dataset using MMseqs2 clustering at various sequence identity thresholds. The left panel illustrates the variation in the total number of clusters as the sequence identity threshold increases from 0.1 to 0.9. The right panel presents a detailed distribution of the number of protein chains per cluster.

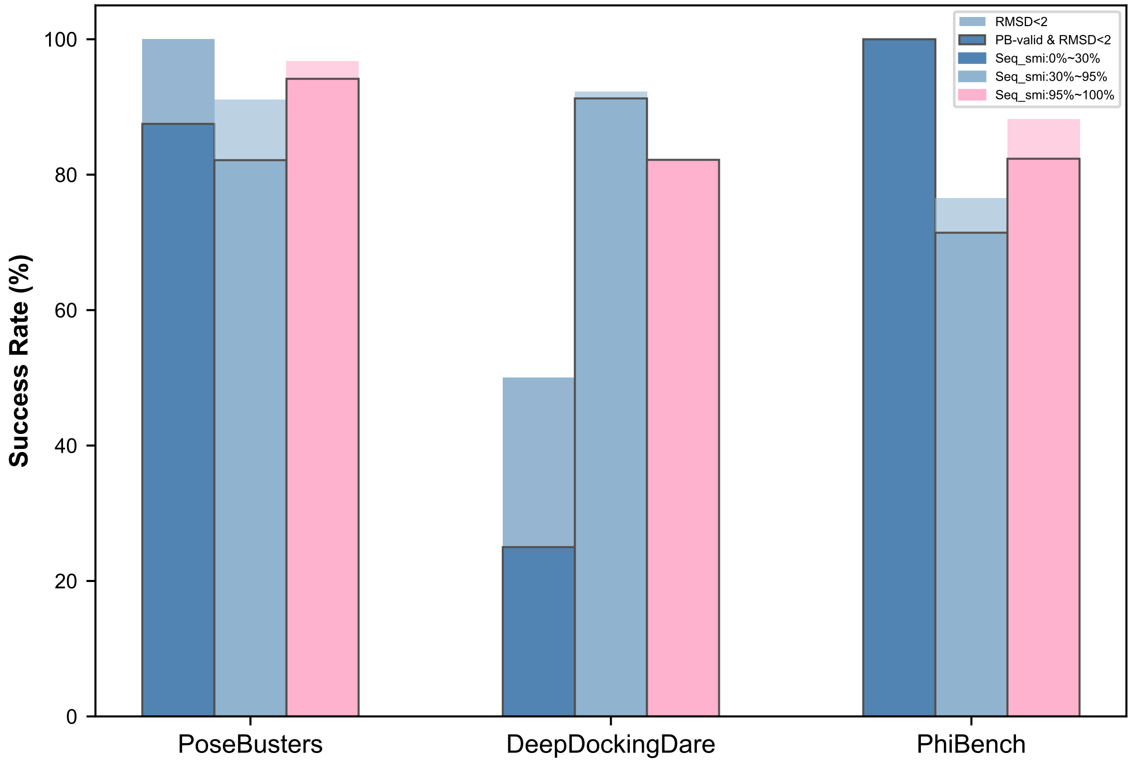

**Figure 16.** Performance of PhysDock on three datasets, categorized by sequence similarity to the Plinder training set (PoseBusters: 428 samples, subsets: 8/ 112/ 308 for low/ medium/ high similarity; DeepDockingDare: 356 samples, subsets: 4/ 206/ 146; PhiBench: 206 samples, subsets: 6/ 98/ 102). Docking success rates are computed under two criteria: PAL-RMSD $\leq$ 2.0 Å, and PB-valid & PAL-RMSD $\leq$ 2.0 Å.

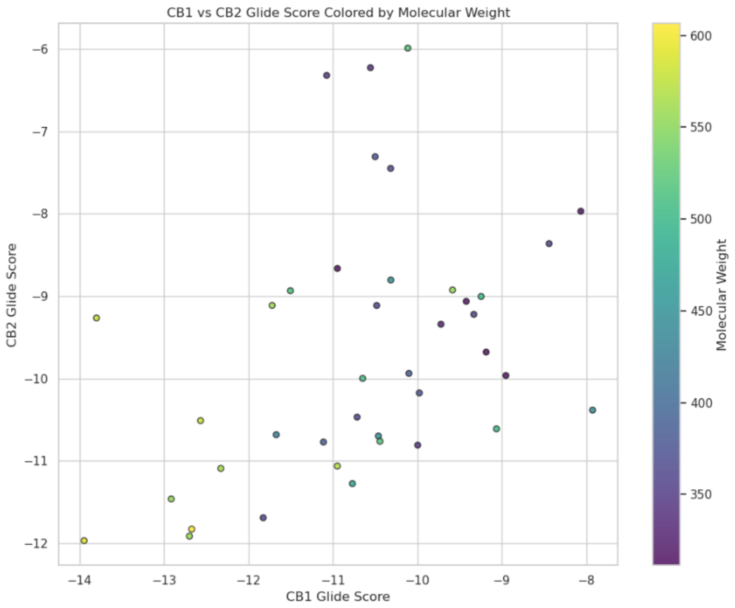

**Figure 17.** Glide scores for 40 compounds docked to CB1 and CB2 receptors using PhysDock. Each point represents a ligand, color-coded by molecular weight.

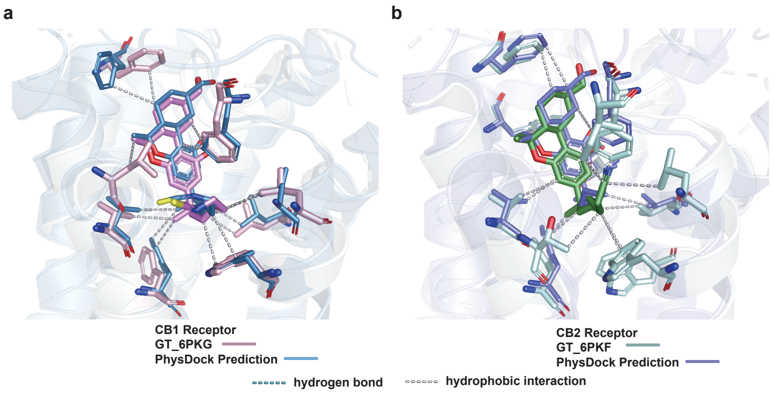

**Figure 18.** The combined pattern graph predicted by Physdock and the overlapping graph of Ground Truth.

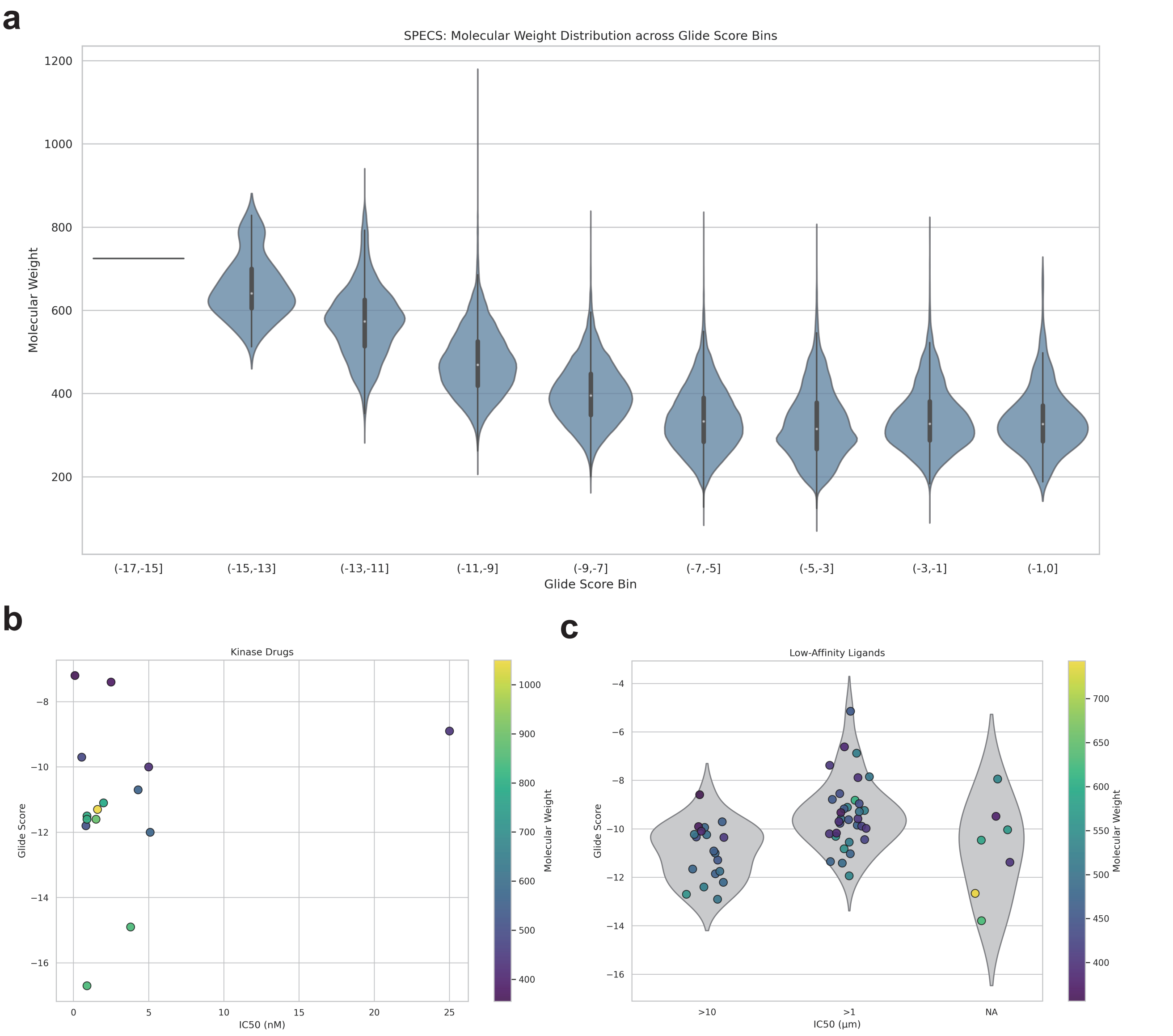

**Figure 19.** **(a)** Violin plot showing the distribution of molecular weights for SPECS compounds across different Glide score bins from PhysDock screening against the NTRK3 kinase. **(b)** Glide scores of known drug candidates docked to the NTRK3 kinase using PhysDock plotted against their $IC_{50}$ values, with points color-coded by molecular weight. **(c)** Glide scores of low-affinity ligands docked to the NTRK3 kinase using PhysDock grouped by $IC_{50}$ categories, with points color-coded by molecular weight.

#### 7 Supplementary Tables

**Table 2.** Docking success rates of existing methods with prior binding pocket information across three benchmark datasets.

| Method | PoseBusters | | DeepDockingDare | | PhiBench | |
|---|---|---|---|---|---|---|
| | PAL-RMSD | PB-valid | PAL-RMSD | PB-valid | PAL-RMSD | PB-valid |
| PhysDock | 408/428 | 389/428 | 317/356 | 309/356 | 171/206 | 160/206 |
| SurfDock | 347/428 | 337/428 | 291/355 | 242/355 | 132/184 | 131/184 |
| Interformer | 332/428 | 300/428 | 251/356 | 229/356 | 123/180 | 114/180 |
| AlphaFold3 | 299/426 | 297/426 | 212/356 | 202/356 | 105/206 | 98/206 |
| Uni-Mol Docking V2 | 257/428 | 209/428 | 142/356 | 135/356 | 98/184 | 96/184 |
| Chai-1 | 267/428 | 264/428 | 140/356 | 127/356 | 92/206 | 87/206 |
| DiffDock-L | 232/428 | 150/428 | 81/344 | 68/344 | 74/186 | 66/186 |
| Glide | 228/428 | 199/428 | 142/356 | 120/356 | 47/185 | 45/185 |
| Vina | 179/428 | 172/428 | 122/355 | 119/355 | 47/198 | 45/198 |
| DiffDock | 186/427 | 104/427 | 75/344 | 66/344 | 57/186 | 48/186 |
| DynamicBind | 188/420 | 185/420 | 82/334 | 81/334 | 100/184 | 98/184 |
| NeuralPLexer | 84/419 | 57/419 | 15/333 | 11/333 | 37/186 | 31/186 |

**Table 3.** Interaction recovery rates and ground-truth interaction counts across three benchmark datasets.

#### PoseBusters

| Model | hydrogen bonds | hydrophobic interactions | salt bridges | pi-stacking | pi-cation interactions |
|---|---|---|---|---|---|
| PhysDock* | 72.2 | 75.6 | 87.6 | 64.0 | 43.1 |
| Uni-Mol Docking V2* | 66.8 | 56.4 | 79.9 | 68.2 | 66.7 |
| SurfDock* | 64.7 | 60.3 | 80.4 | 50.6 | 55.0 |
| Glide* | 51.0 | 56.2 | 75.8 | 44.3 | 26.9 |
| DiffDock-l* | 54.9 | 43.8 | 67.7 | 50.7 | 32.5 |
| DynamicBind | 56.2 | 52.3 | 76.4 | 38.7 | 21.2 |
| Interformer* | 46.1 | 45.0 | 56.5 | 53.6 | 0.0 |
| Vina* | 39.7 | 41.0 | 49.3 | 40.9 | 19.5 |
| DiffDock* | 41.8 | 37.1 | 52.2 | 36.7 | 22.5 |
| AlphaFold3 | 48.2 | 37.6 | 17.6 | 11.5 | 10.4 |
| Chai-1 | 44.9 | 26.1 | 17.7 | 8.0 | 6.9 |
| NeuralPLexer | 16.6 | 14.0 | 20.7 | 11.4 | 8.5 |

**Interaction counts — Hydrogen bonds: 1688, Hydrophobic interactions: 1323, Salt bridges: 369, Pi-stacking: 170, Pi-cation interactions: 52**

#### DeepDockingDare

| Model | hydrogen bonds | hydrophobic interactions | salt bridges | pi-stacking | pi-cation interactions |
|---|---|---|---|---|---|
| Uni-Mol Docking V2* | 56.3 | 52.1 | 83.2 | 54.5 | 62.9 |
| SurfDock* | 56.9 | 56.3 | 78.4 | 38.8 | 46.3 |
| PhysDock* | 43.0 | 59.3 | 38.7 | 44.9 | 23.5 |
| Interformer* | 34.5 | 40.8 | 48.5 | 42.1 | 41.7 |
| Vina* | 30.3 | 36.7 | 48.0 | 42.4 | 38.8 |
| Glide* | 33.7 | 39.5 | 33.7 | 37.8 | 14.3 |
| DiffDock-l* | 30.7 | 25.6 | 48.9 | 23.2 | 26.6 |
| DiffDock* | 25.8 | 23.7 | 47.7 | 23.3 | 26.5 |
| DynamicBind | 30.3 | 22.9 | 53.2 | 17.1 | 14.8 |
| AlphaFold3 | 30.8 | 24.0 | 31.5 | 32.1 | 13.6 |
| Chai-1 | 27.7 | 15.4 | 22.4 | 10.7 | 7.8 |
| NeuralPLexer | 9.4 | 12.4 | 9.3 | 16.2 | 1.1 |

**Interaction counts — Hydrogen bonds: 893, Hydrophobic interactions: 1548, Salt bridges: 97, Pi-stacking: 41, Pi-cation interactions: 23**

#### PhiBench

| Model | hydrogen bonds | hydrophobic interactions | salt bridges | pi-stacking | pi-cation interactions |
|---|---|---|---|---|---|
| Uni-Mol Docking V2* | 63.4 | 49.0 | 85.9 | 63.4 | 50.8 |
| SurfDock* | 57.2 | 48.7 | 82.7 | 35.1 | 38.0 |
| PhysDock* | 52.6 | 53.5 | 51.9 | 58.2 | 17.8 |
| DiffDock-l* | 53.4 | 37.6 | 73.6 | 34.9 | 29.5 |
| Glide* | 43.0 | 42.7 | 79.2 | 31.8 | 17.4 |
| Interformer* | 37.8 | 36.0 | 51.1 | 43.6 | 28.2 |
| DynamicBind | 47.5 | 36.8 | 57.4 | 29.6 | 14.9 |
| DiffDock* | 39.7 | 32.5 | 62.4 | 26.2 | 17.6 |
| Vina* | 34.7 | 31.6 | 60.1 | 21.7 | 24.2 |
| Chai-1 | 46.3 | 18.9 | 20.2 | 11.5 | 6.8 |
| NeuralPLexer | 27.1 | 16.7 | 30.6 | 9.0 | 12.9 |
| AlphaFold3 | 45.2 | 30.0 | 14.1 | 0.0 | 5.0 |

**Interaction counts — Hydrogen bonds: 1002, Hydrophobic interactions: 583, Salt bridges: 164, Pi-stacking: 33, Pi-cation interactions: 38**

**Table 4.** Experimental binding free energies and Glide scores for all compounds docked to CB1 and CB2 receptors using PhysDock.

| Name | Molecules | CB1_ΔG_exp (kcal/mol) | CB1_PhysDock Glide (kcal/mol) | CB2_ΔG_exp (kcal/mol) | CB2_PhysDock Glide (kcal/mol) |
|---|---|---|---|---|---|
| MK-0364 [2] | 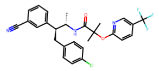 | -13.49 | -11.501 | -9.23 | -8.935 |
| ACEA [1] | 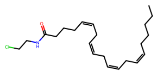 | -12.15 | -10.315 | -7.82 | -7.45 |
| AM-11542 [2] | 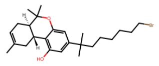 | -12.12 | -10.46 | -13.59 | -10.699 |
| ACPA [1] | 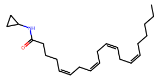 | -11.88 | -10.554 | -8.44 | -6.228 |
| THC [2] | 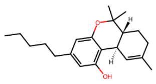 | -11.65 | -9.42 | -10.08 | -9.064 |
| SR-147778 [2] | 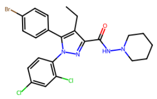 | -11.53 | -10.111 | -8.73 | -5.99 |
| AM-251 [2] | 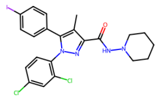 | -11.08 | -9.582 | -7.69 | -8.926 |
| WIN-55,212-2 [2] | 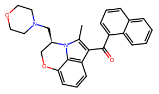 | -10.96 | -11.669 | -11.84 | -10.683 |
| R-methanandamide [1] | 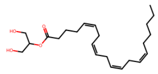 | -10.57 | -10.48 | -8.36 | -9.114 |
| 2-AG [1] | 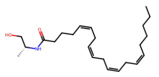 | -9.93 | -10.498 | -9.39 | -7.308 |

| Name | Molecules | CB1_ΔG_exp (kcal/mol) | CB1_PhysDock Glide (kcal/mol) | CB2_ΔG_exp (kcal/mol) | CB2_PhysDock Glide (kcal/mol) |
|---|---|---|---|---|---|
| anandamide [1] | 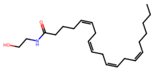 | -9.9 | -11.072 | -7.84 | -6.321 |
| PNR-4-20 [1] | 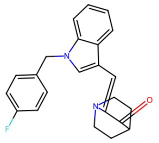 | -9.33 | -11.823 | -12 | -11.691 |
| UR-144 [2] | 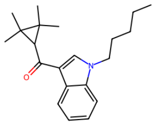 | -9.3 | -10.945 | -11.93 | -8.665 |
| SR-144528 [2] | 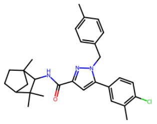 | -8.73 | -10.767 | -12.58 | -11.277 |
| VJ-115 [1] | 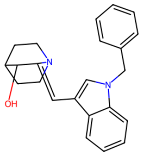 | -7.52 | -9.995 | -8.6 | -10.809 |
| CBD [2] | 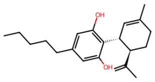 | -7.24 | -8.063 | -7.334 | -7.97 |
| AM-630 [2] | 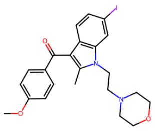 | -7.2 | -9.244 | -10.23 | -9.005 |
| GW842166X [3] | 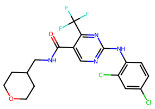 | >-6.17 | -10.313 | -9.96 | -8.805 |
| JWH-151 [3] | 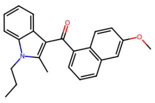 | >-6.82 | -10.712 | -10.26 | -10.469 |
| L-759,633 [4] | 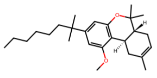 | -6.55 | -10.098 | -10.5 | -9.939 |

| Name | Molecules | CB1_ΔG_exp (kcal/mol) | CB1_PhysDock Glide (kcal/mol) | CB2_ΔG_exp (kcal/mol) | CB2_PhysDock Glide (kcal/mol) |
|---|---|---|---|---|---|
| GW-405,833 [5] | 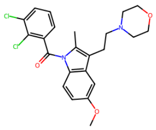 | -7.26 | -7.924 | -11.49 | -10.384 |
| PF-03550096 [5] | 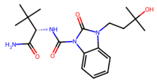 | -11.05 | -9.975 | -7.96 | -10.176 |
| APD371 [5] | 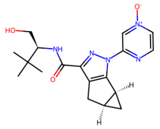 | >-6.82 | -9.33 | -11.2 | -9.222 |
| Cannabilactone [5] | 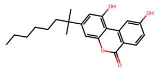 | -8.73 | -8.439 | -12.4 | -8.363 |
| HU-211 [5] | 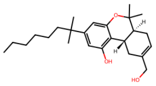 | -7.78 | -11.11 | >-6.82 | -10.772 |
| 7 [6] | 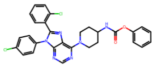 | -11.72 | -11.718 | -6.78 | -9.113 |
| 9 [6] | 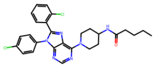 | -10.52 | -10.441 | -8.16 | -10.763 |
| 10 [6] | 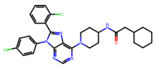 | -11.89 | -12.325 | -7.68 | -11.092 |
| 18 [6] | 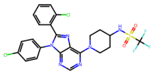 | -11.21 | -10.946 | -7.29 | -11.063 |
| Otenabant [7] | 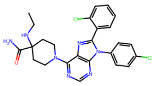 | -12.49 | -10.646 | -6.98 | -9.999 |

| Name | Molecules | CB1_ΔG_exp (kcal/mol) | CB1_PhysDock Glide (kcal/mol) | CB2_ΔG_exp (kcal/mol) | CB2_PhysDock Glide (kcal/mol) |
|---|---|---|---|---|---|
| 12 [7] | 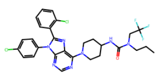 | -12.22 | -12.67 | -7.58 | -11.8287 |
| 15 [7] | 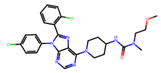 | -13.08 | -12.912 | -7.69 | -11.463 |
| 25 [7] | 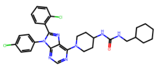 | -12.34 | -12.565 | -6.98 | -10.512 |
| 29 [7] | 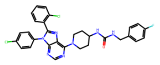 | -11.63 | -13.944 | >-6.82 | -11.969 |
| 30 [7] | 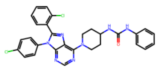 | -11.12 | -12.697 | >-6.82 | -11.914 |
| 32 [7] | 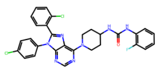 | -11.46 | -13.797 | >-6.82 | -9.266 |
| AM1241 [8] | 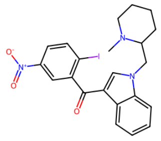 | -7.23 | -9.062 | -10.67 | -10.612 |
| 24 [9] | 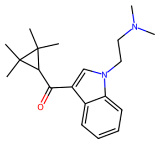 | >-6.82 | -9.184 | -11.9 | -9.68 |
| 36 [9] | 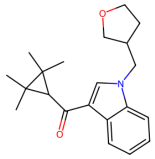 | -8.46 | -9.719 | -13.44 | -9.342 |
| 50 [9] | 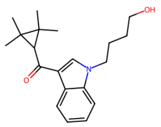 | >-7.16 | -8.95 | -11.81 | -9.964 |

**Table 5.** Docking results of kinase inhibitors across different models

| Name | Molecules | Stage | IC50 (nM) | PhysDock_Glide (kcal/mol) |
|---|---|---|---|---|
| Repotrectinib | 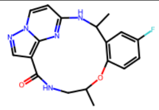 | Approved | 0.1 | -7.2 |
| Entrectinib | 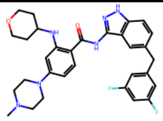 | Approved | 4.3 | -10.7 |
| Larotrectinib | 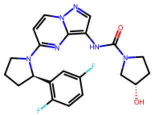 | Approved | 5-11 | -10.0 |
| Lestaurtinib | 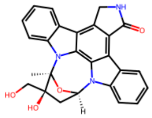 | Phase III | 25 | -8.9 |
| Selitrectinib | 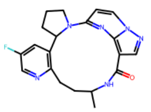 | Phase I | 2.5 | -7.4 |
| CH7057288 | 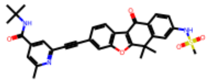 | Preclinical | 5.1 | -12.0 |
| Altiratinib | 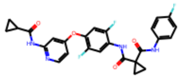 | Preclinical | 0.83 | -11.8 |
| CPD-085 | 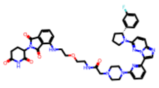 | Preclinical | 3.8 | -14.9 |
| Compound 30f | 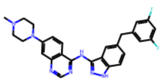 | Preclinical | 0.55 | -9.7 |
| CPD-141 | 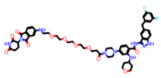 | Preclinical | 1.6 | -11.3 |
| CPD-145 | 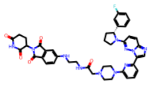 | Preclinical | 0.9 | -11.5 |
| CPD-053 | 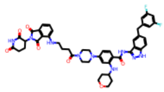 | Preclinical | 1.5 | -11.6 |
| CPD-211 | 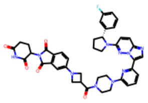 | Preclinical | 2 | -11.1 |
| CPD-143 | 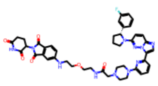 | No progress | 0.9 | -16.7 |
| CPD-159 | 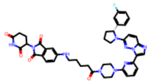 | No progress | 0.9 | -11.6 |

**Table 6.** Experimental $IC_{50}$ values and Glide scores for known drug candidates and weak binders docked to NTRK3 kinase using PhysDock.

| Name | Molecules | IC50 (µM) | PhysDock_Glide (kcal/mol) |
|---|---|---|---|
| B30 [10] | 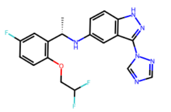 | NA | -11.4 |
| 7n [11] | 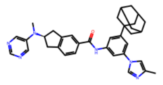 | >10 | -12.7 |
| 7m [11] | 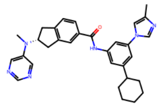 | >10 | -12.9 |
| 7l [11] | 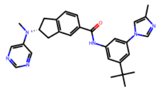 | >10 | -12.2 |
| 7k [11] | 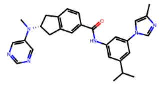 | >10 | -11.9 |
| 7j [11] | 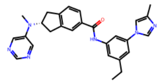 | >10 | -11.3 |
| 7i [11] | 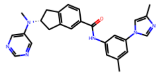 | >10 | -10.3 |
| 7h [11] | 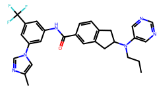 | >10 | -12.4 |
| 7g [11] | 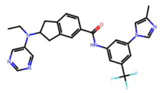 | >10 | -10.2 |
| 7f [11] | 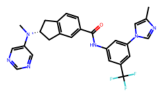 | >10 | -11.8 |

| Name | Molecules | IC50 (μM) | PhysDock_Glide (kcal/mol) |
|---|---|---|---|
| 7e [11] | 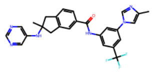 | >10 | -10.2 |
| 7d [11] | 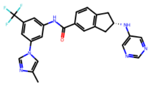 | >10 | -11.0 |
| 7c [11] | 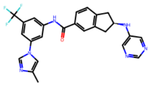 | >10 | -11.7 |
| 7b [11] | 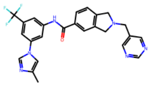 | >10 | -9.9 |
| 7a [11] | 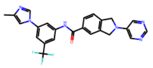 | >10 | -9.7 |
| 6n [12] | 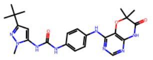 | >10 | -10.9 |
| 6k [12] | 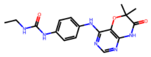 | >10 | -8.6 |
| 6j [12] | 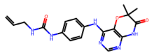 | >10 | -9.9 |
| 6i [12] | 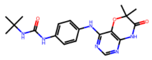 | >10 | -10.1 |
| 6g [12] | 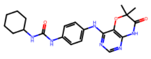 | >10 | -10.4 |

| Name | Molecules | IC50 (μM) | PhysDock_Glide (kcal/mol) |
|---|---|---|---|
| 32h [13] | 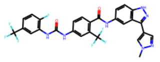 | >1 | -8.8 |
| 32e [13] | 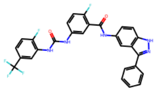 | >1 | -10.8 |
| 32c [13] | 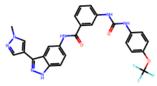 | >1 | -11.9 |
| 32b [13] | 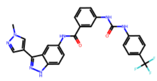 | >1 | -6.9 |
| 32a [13] | 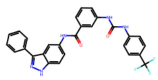 | >1 | -10.5 |
| 26k [13] | 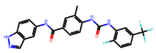 | >1 | -9.8 |
| 26h [13] | 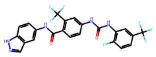 | >1 | -9.1 |
| 26g [13] | 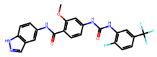 | >1 | -11.4 |
| 26c [13] | 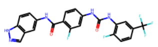 | >1 | -11.4 |
| 26a [13] | 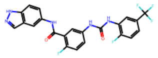 | >1 | -11.0 |

| Name | Molecules | IC50 (µM) | PhysDock_Glide (kcal/mol) |
|---|---|---|---|
| 23c [13] | 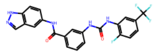 | >1 | -10.4 |
| 23b [13] | 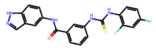 | >1 | -9.2 |
| 23a [13] | 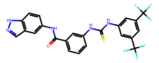 | >1 | -5.1 |
| 21b [13] | 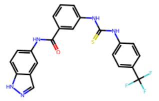 | >1 | -8.5 |
| 21a [13] | 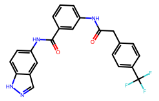 | >1 | -9.8 |
| 19c [13] | 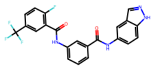 | >1 | -10.3 |
| 19b [13] | 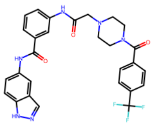 | >1 | -9.6 |
| 16z [13] | 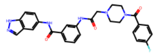 | >1 | -9.0 |
| 16y [13] | 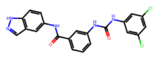 | >1 | -7.8 |
| 16w [13] | 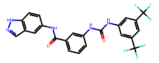 | >1 | -8.8 |

| Name | Molecules | IC50 (μM) | PhysDock_Glide (kcal/mol) |
|---|---|---|---|
| 16v [13] | 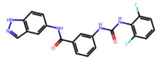 | >1 | -7.4 |
| 16t [13] | 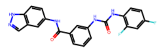 | >1 | -9.6 |
| 16s [13] | 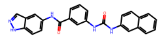 | >1 | -9.9 |
| 16o [13] | 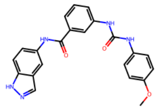 | >1 | -9.7 |
| 16m [13] | 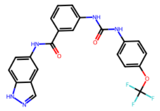 | >1 | -9.2 |
| 16i [13] | 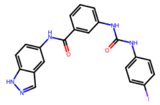 | >1 | -9.3 |
| 16h [13] | 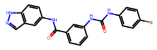 | >1 | -9.6 |
| 16g [13] | 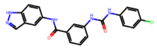 | >1 | -10.0 |
| 16f [13] | 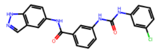 | >1 | -10.2 |
| 16d [13] | 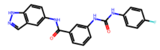 | >1 | -10.2 |

| Name | Molecules | IC50 (µM) | PhysDock_Glide (kcal/mol) |
|---|---|---|---|
| 16c [13] | 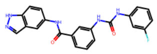 | >1 | -6.6 |
| 16aa [13] | 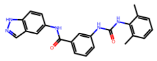 | >1 | -7.9 |
| 16a [13] | 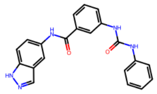 | >1 | -9.3 |
| 22 [14] | 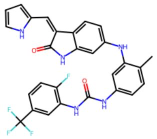 | NA | -7.9 |
| 19 [14] | 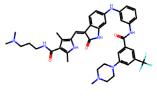 | NA | -12.7 |
| 16 [14] | 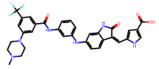 | NA | -13.8 |
| 7 [14] | 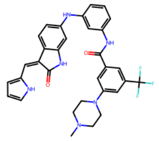 | NA | -10.5 |
| 5_1 [15] | 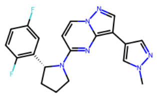 | NA | -9.5 |
| 5_2 [14] | 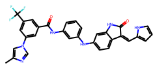 | NA | -10.0 |
